# Seasonal dynamics in the trophic ecology and condition of a marine, benthic mesopredator, the southern stingray, *Hypanus americanus*

**DOI:** 10.64898/2026.02.02.703287

**Authors:** Vital Heim, Matthew J. Smukall, Natalie D. Mylniczenko, Charlene M. Burns, Nigel E. Hussey, Ansgar Kahmen, Philip Matich

**Affiliations:** Department of Environmental Sciences, Zoology, University of Basel, Vesalgasse 1, 4051 Basel, Switzerland; Bimini Biological Field Station Foundation, South Bimini, The Bahamas; Department of Animal Health, Disney’s Animals, Science & Environment, Animal Programs, 1200 Savannah Cir, Lake Buena Vista, FL 32830, U.S.A.; Department of Integrative Biology, University of Windsor, 401 Sunset Avenue, Windsor, Ontario N9B 3P4, Canada; Department of Environmental Sciences, Botany, University of Basel, Schönbeinstrasse 6, 4056 Basel, Switzerland; Saving The Blue, Cooper City, FL 33328, U.S.A.; Department of Marine Biology, Texas A&M University at Galveston, Galveston, Texas 77553, U.S.A.

**Keywords:** Bayesian mixing models, chemical tracers, elasmobranch, mesopredator release nutritional state

## Abstract

Mesopredators contribute to food web stability and as such, understanding their trophic ecology can help to predict potential consequences of ongoing ecosystem modification. Here, multi-tissue carbon and nitrogen stable isotope analysis (δ^13^C and δ^15^N) and biochemical blood parameters (β-hydroxybutyrate, glucose, lactate, and osmolality) were used to assess sex, size, spatial and seasonal differences in trophic ecology and condition of southern stingrays, *Hypanus americanus,* in Bimini, The Bahamas. Stingrays exhibited a dietary preference for molluscs and annelids, with an ontogenetic shift towards lower δ^13^C with increasing body size indicating a shift towards more mangrove associated prey. Nitrogen isotope values showed minimal seasonal changes, but higher δ^15^N values in males indicated foraging at a higher trophic level than females. Blood β-hydroxybutyrate concentrations and osmolality revealed a similar energetic state and condition between sex, size, location and season. Our results advance our understanding of the seasonal trophic ecology of a benthic, marine mesopredator and identify the southern stingray as an important trophic link in seagrass and mangrove habitats in Bimini.

## 1. Introduction

The global loss of biodiversity and removal of large predators is altering the structure of food webs, with ramifications for ecosystem function and how ecological systems will respond to continued perturbations (Estes et al., 2011; McCauley et al., 2015). Understanding the trophic ecology of individual species is therefore crucial for wildlife conservation and the preservation of ecosystem stability (Pace et al., 1999). Apex predators have been the focus of several studies describing their ecological role in maintaining ecosystem dynamics through top-down regulation of lower trophic levels via direct (i.e. predation), and indirect (i.e. risk) effects (Heithaus et al., 2007; Ripple & Beschta, 2004; Ritchie & Johnson, 2009). How their removal jeopardises ecosystem regulation in part depends on the mediation of trophic links by species at intermediate trophic positions (i.e. mesopredators), which play an integral part in stabilising food webs (Bascompte et al., 2005; Fagan, 1997; Ritchie & Johnson, 2009, Shipley et al., 2023). In marine food webs, these regulatory functions are often performed by elasmobranchs (i.e. sharks and rays; Flowers et al., 2021; Heithaus et al., 2010; Heupel et al., 2014), and in light of the declines of apex predatory sharks, there is growing concern for the risk of mesopredator release and subsequent trophic cascades in these environments (Burkholder et al., 2013; Ferretti et al., 2010; Heithaus et al., 2012; Pershing et al., 2015; Pinnegar et al., 2000). Collecting the necessary data to advance our understanding of these risks remains difficult due to logistic challenges in observing marine food webs, their complexity, concealing environments and a lack of observation time. Consequently, the functional roles of marine mesopredators often remain underappreciated limiting our capability to predict the outcome and intensity of potential cascades on specific trophic levels, and they therefore require greater attention as we seek to improve wildlife management (Burt et al., 2018; Grubbs et al., 2016).

Many rays of the superorder Batoidea often act as mesopredators of benthic infauna and other invertebrates in marine ecosystems (Flowers et al., 2021; O’Shea et al., 2018). Batoid rays can be quite abundant, and are prey to larger marine predators, including sharks, killer whales, *Orcinus orca*, and other rays (Higuera-Rivas et al., 2023; Poulakis et al., 2017; Raoult et al., 2019). Batoidea impact the density and diverstiy of lower trophic levels through foraging (Glaspie & Seitz, 2017), can facilitate foraging symbioses with other species (Kajiura et al., 2009, Kiszka et al., 2015) and given their mobility, link different ecosystem components (Earl & Zollner, 2017; Papastamatiou et al., 2015). Furthermore, rays are important bioturbators redistributing nutrients in the water column that can stimulate productivity (O’Shea et al., 2012; Takeuchi & Tamaki, 2014). While significant research has examined the overall ecology of rays (reviewed by Flowers et al., 2021), detailed, species-specific insights into their trophic ecology would improve understanding of their importance for ecosystem functioning and stability.

The trophic ecology and role of a species within a food web can be influenced by a variety of factors such as seasonal variability in environmental conditions (McMeans et al., 2019; Rogers et al., 2020), ontogenetic dietary shifts (Matich et al., 2019; Sánchez-Hernández et al., 2022), predation risk (Heithaus et al., 2009), and habitat availability (Hayden et al., 2019). An additional factor known to be connected to the ecological role of an animal is its condition (Barley et al., 2017). The condition of an animal, often discussed in terms of relative energy stores (Green 2001, Schick et al., 2013), can vary in response to habitat use and variables determining habitat quality such as nutrient availability, temperature, reproduction, and environmental stressors (Hussey et al., 2009; Lyons et al., 2017; Rangel et al., 2022; Weideli et al., 2019). An animal’s condition can also be impacted by the presence of predators, either directly through predation (e.g. physical injury or death; Ritchie & Johnson, 2009), or indirectly through energetically costly stress responses associated with predation risk such as inhabiting unprofitable environments (Pérez-Tris et al., 2004; Preisser et al., 2005). Understanding how the condition of an animal changes through time and space can not only reveal how successful wild animals are at securing resources but also provides insights into population and food web dynamics (Heithaus et al., 2012; Schick et al., 2013).

Assessing variation in the condition of wild elasmobranchs remains challenging, with several approaches adopted. The liver is the main storage site of lipids in elasmobranchs and a lipid-rich, large liver can indicate an elevated nutritional state (i.e. improved condition with increasing hepatosomatic index (HSI); Ballantyne, 1997; Watson & Dickson, 2001; Zammit & Newsholme, 1979). Consequently, a variety of condition indices using liver mass, or approximations thereof, i.e. weight-to-length or span-to-length relationships (i.e. Fulton condition factor [CF]), have been used to assess the energetic and nutritional state of individuals, but with mixed results (Gallagher, Wagner, et al., 2014; Hussey et al., 2009; Irschick & Hammerschlag, 2014; Wheeler et al., 2023). Drawbacks of these methods include lethal sampling (for HSI) and limited sensitivity (CF) (Hussey et al., 2009), and therefore alternative methods are needed. With advancements in technology, ultrasonography has proven a reliable method to estimate liver size, mass and density in aquarium-managed elasmobranchs (Grant et al., 2013; Mylniczenko, 2012; Mylniczenko et al., 2017). Yet, there is a need for a non-lethal alternative that does not require expensive ultrasonography (e.g. chemical tracers; Bouyoucos et al., 2017; Hammerschlag et al., 2018; Moorhead et al., 2021; Prohaska et al., 2021; Talwar et al., 2017).

Sharks and rays have very little adipose tissue and a limited capacity to oxidize storage lipids outside of the liver, thus they rely on lipid-derived ketone bodies to fuel oxidative processes in extrahepatic tissues (e.g. skeletal muscle) or as a main energy source during starvation (Ballantyne, 1997, 2015, 2016; deRoos, 1994; Speers-Roesch & Treberg, 2010). Therefore, elevated concentrations of ketone bodies such as β-hydroxybutyrate in blood samples result from physical activity (Ballantyne, 1997, Valls et al., 2016), chronic stress such as fasting and long-term anorexia (Wood et al., 2010; Zammit & Newsholme, 1979), and subsequently lower condition (deRoos et al., 1985; Moorhead et al., 2021). In response to an acute health impact, the concentrations of glucose and lactate in blood are expected to increase in elasmobranchs (Gallagher, Serafy, et al., 2014; Skomal, 2007; Skomal & Mandelman, 2012). Elevated glucose levels indicate increased mobilisation of glycogen storages in the liver through secretion of catecholamines and corticosteroids to support increased brain and muscle needs (Cliff & Thurman, 1984; deRoos, 1994; deRoos & deRoos, 1978; Hoffmayer & Parsons, 2001; Marshall et al., 2012; Skomal & Mandelman, 2012). An increase in lactate indicates muscle exertion, anaerobic respiration and hypoventilation (Knotek et al., 2022; Moyes et al., 2006; Prohaska et al., 2018; Talwar et al., 2017). In combination, lactate and glucose can be used to tease apart the causes of a stress response. For example, both, glucose and lactate concentrations were elevated in juvenile lemon sharks, *Negaprion brevirostris*, after simulated capture on a longline (Bouyoucos et al., 2017), and in Atlantic stingrays, *Hypanus sabinus,* when they were temporarily removed from the water (Lambert et al., 2018). Urea is synthesised in the liver and changes in blood urea nitrogen are a good indicator of animal condition as blood urea nitrogen reduces with decreasing liver size. Due to high urea concentrations in their body fluids, elasmobranchs are expected to always be hyper- or at least iso-osmolar compared to the surrounding seawater (Ballantyne, 1997, 2015; Deck et al., 2016). Changes in blood plasma osmolality can therefore be used to diagnose a fluid shift resulting from decreasing urea levels as a result of lower liver condition following sustained and severe stress (e.g. chronic starvation; Armour et al., 1993; Ballantyne, 2015; Fuller et al., 2020; Skomal & Bernal, 2010). Together, these four metrics provide a snapshot of the metabolic state, hereafter referred to as condition, of an individual in lieu of physical variables or expensive equipment (e.g. body or liver weight, ultrasound). However, species-specific differences and a lack of baseline values and comparison between these four metrics, challenge our ability to reliably use these tracers to assess elasmobranch condition in the wild.

Here, we assess the trophic ecology and condition of a benthic, marine elasmobranch mesopredator, the southern stingray, *Hypanus americanus* (Hildebrand & Schroeder, 1928), in Bimini, The Bahamas. Southern stingrays are found in shallow estuaries, sand, seagrass, and coral reef habitats down to ∼ 100 m depth in the Western Atlantic from New Jersey (U.S.A.) to Brazil including The Bahamas and Caribbean (Carlson et al., 2020; Last et al., 2016). Despite being one of the most abundant rays in the Western Atlantic, relatively little information exists regarding their trophic ecology (O’Shea et al., 2020). As mesopredators, southern stingrays are expected to be subjected to top-down control from larger, apex predators such as great hammerheads, *Sphyrna mokarran* (Cliff, 1995; Guttridge et al., 2025; Roemer et al., 2016; Strong et al., 1990), while they regulate lower trophic levels via direct and indirect predator effects (Flowers et al., 2021; Sherman et al., 2020). Consequently, southern stingrays are hypothesised to provide an important connection between lower and upper trophic levels in coastal food webs (Ritchie & Johnson, 2009) and are a keystone species in nearshore marine ecosystems (O’Shea et al., 2012, 2020). Available stomach content analyses indicate dietary preferences for crustaceans, teleosts, annelids and molluscs (Bigelow & Schroeder, 1953; Bowman et al., 2000; Gilliam & Sullivan, 1993; O’Shea et al., 2020; Randall, 1967; Smith & Herrnkind, 1992; Tilley et al., 2013), which is supported by initial studies using chemical tracer methods (i.e. stable isotopes and fatty acids) to elucidate the broader trophic interactions of southern stingrays (Hoopes et al., 2020; O’Shea et al., 2020; Shipley et al., 2018; Tilley et al., 2013). Here, we used carbon and nitrogen stable isotopes (δ^13^C and δ^15^N) in white muscle tissue and blood to undertake a comprehensive assessment of their trophic ecology when considering sex, size, capture location and season. We then examine variation in concentrations of β-hydroxybutyrate, glucose, lactate and osmolality in blood samples to provide a baseline for those four condition metrics in wild caught southern stingrays.

## 2. Material and Methods

### 2.1. Study site

This study was conducted in the waters surrounding Bimini, The Bahamas (25° 44’ N, 79° 15’ W), a group of two small mangrove-fringed islands that lie at the western edge of the Great Bahama Bank approximately 86 km east of Miami (Florida, U.S.A., Fig. 1).

**Fig. 1.**
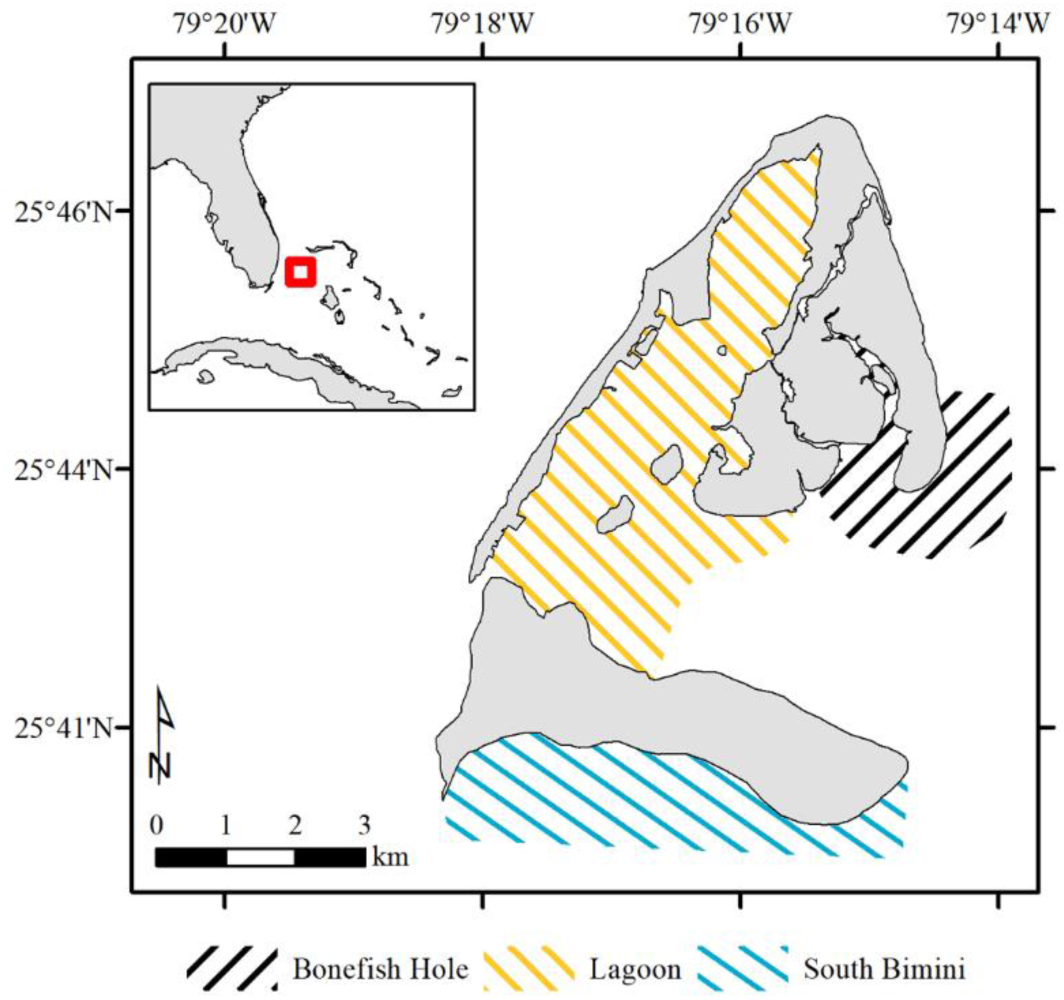
Study site and capture areas. The insert in the top left shows the location of the Bimini Islands in the western North Atlantic. The three different capture locations are shown in different colours for Bonefish Hole (black), the lagoon (yellow), and South Bimini (cyan). Most southern stingray captures (N = 130) were recorded in the South Bimini area, followed by the lagoon and the Bonefish Hole area (N = 24, each).

The islands and their surroundings offer a wide range of habitat types including a shallow-water lagoon between South and North Bimini, shallow-water sand flats, seagrass, and mixed sponge and rocky hard bottom habitats to the east and south, and fringing coral reefs to the west that extend into the shelf habitat and the deep waters of the Gulf Stream (Jennings et al., 2012; van Zinnicq Bergmann et al., 2022). Southern stingrays are present throughout the study site year-round (van Zinnicq Bergmann et al., 2022). However, there is higher residency of apex predatory sharks such as great hammerheads and tiger sharks, *Galeocerdo cuvier,* during the cooler water winter months (Guttridge et al., 2017; Heim et al., 2021, 2024, Smukall et al., 2022), and therefore predation risk to southern stingrays is expected to replicate this seasonal pattern. The three capture locations (Fig. 1) differ not only in microhabitat type (see van Zinnicq Bergmann et al., 2024), but also in predator exposure, with the lagoon becoming inaccessible to larger predators during low tide (Guttridge et al., 2012). Previous work in this system (see van Zinnicq Bergmann et al., 2024) revealed that southern stingrays preferentially select the lagoon for safety over resource availability.

### 2.2. Southern stingray capture and tissue sample collection

Southern stingrays were caught year-round in shallow waters (< 1.5 m) between July 10^th^, 2018, and November 27^th^, 2019. Two flat-bottom skiffs (16 and 17 ft) were used to visually locate rays and then follow or guide them towards the mangroves or beach upon sight. Once the southern stingrays were in a shallow enough area, the skiff crews encircled an individual using panels (∼ 1×1 m) of fencing mesh attached to two PVC poles, and large dip nets (∼ 80 x 90 cm, 15/16” square knotless nylon, Loki Nets, Knoxville, Tennessee, USA). Once encircled, the southern stingrays were caught using the dip nets and immediately transferred into a seawater-filled tub (∼ 150 cm diameter, ∼ 50 cm height) on board the skiffs. The duration from initially spotting a southern stingray to its successful capture varied considerably from approximately one minute to ∼ 30 minutes, but no specific capture time was recorded. Depending on the capture location, southern stingrays were measured and sampled directly on the boat (Lagoon and Bonefish Hole, Fig. 1) or transported to a semi-captive pen, to be measured and sampled the following day (South Bimini, Fig. 1).

Once a southern stingray was placed in the tub, it was gently restrained by holding its pectoral fins, and the venomous barb of the ray was immediately wrapped with a thick piece of fabric. All southern stingrays were scanned for the presence of a passive integrated transponder (PIT, Biomark, Boise, Idaho, U.S.A.) indicating the individual had been previously caught. In the absence of a PIT tag (i.e. a newly caught individual), a new one was inserted at the base of the tail using the inserter (MK7 Implanter) provided by the manufacturer. In the rare event that the same individual was recaptured within less than two weeks, it was released immediately. The disc width (i.e. the distance between the two most distal points of the stingrays’ pectoral fins) was recorded to the nearest 0.1 cm. Individuals were sexed based on the presence or absence of claspers (i.e. the external reproductive organs of males). Female individuals were subsequently categorised as young-of-the-year, sexually immature or sexually mature based on published disc-width-at-maturity values (Hayne et al., 2018; Henningsen & Leaf, 2010). Sexual maturity in males was based on clasper calcification (Clark & von Schmidt, 1965).

White muscle tissue samples, hereafter “muscle samples”, were taken from the pectoral fin of the stingrays using a medical muscle biopsy punch (5 – 10 mm diameter, depending on the size of the individual). Blood samples were collected via venipuncture (18 – 23 g x 1.5-inch needles, depending on the size of the animal) of the caudal tail vein after the southern stingrays were restrained in dorsal recumbency. Depending on the size of the sampled individual, 3 – 12 ml blood were collected. In the case of a recapture event, a new blood sample and muscle sample was taken if at least 2 and 4 weeks respectively had elapsed since last capture. In the field, muscle and blood samples were kept in a cooler with ice to prevent tissue degradation. Upon return to the land-based facility, muscle samples were immediately frozen at – 20° C. Blood samples were transferred into non-heparinised 5 ml cryovials and centrifuged (3500 rpm, E8 centrifuge, LW Scientific, Lawrenceville, Georgia, U.S.A.) until separated into blood plasma and red blood cells (∼ 2 – 5 minutes). The use of non-heparinised containers means that, by definition, we did not actually aliquot blood plasma. However, blood samples were centrifuged in an uncoagulated form and thus the liquid state that was later aliquoted contained coagulation proteins (Issaq et al., 2007; Krebs, 1950). For simplicity, we thus use the term “serum” to refer to the liquid, aliquoted part of the centrifuged blood sample hereafter. Serum was aliquoted into non-heparinised 2.0 ml cryovials and frozen at -20° C until analysed. We aliquoted 0.5 ml serum for the stable isotope analysis (see 2.3.), and 1.0 ml serum for the measurement of the condition metrics (see 2.4.). Remaining serum and the red blood cells were stored at -20° C for use in other studies.

### 2.3. Stable isotope analysis

We used stable isotope analysis of carbon and nitrogen isotopes as a minimally invasive method to investigate the trophic ecology of southern stingrays. Muscle and serum samples were chosen based on the different tissue turnover rates of these tissues, which allows for the exploration of isotopic variation across different time scales (Bearhop et al., 2002). Currently there is no species-specific isotope data available on isotopic turnover rates of tissues for southern stingrays. Consequently, we used data from freshwater ocellate river stingrays, *Potamotrygon motoro,* as an approximation: 98 – 422 days in white muscle tissue, and 53 – 265 days in blood samples (MacNeil et al., 2006).

Following the removal of connective tissue using sterilized scissors, the white muscle tissue was rinsed with deionised water and re-frozen at -20° C. Muscle samples were analysed at two different laboratories: (1) at the Movement and Trophic Ecology Laboratory at the University of Windsor, Ontario, Canada (N = 153, Table S1) and (2) at the Stable Isotope Ecology Lab at the University of Basel, Switzerland (N = 85, Table S1). All serum samples were analysed at the University of Basel.

Samples analysed at the University of Windsor were freeze-dried for 48 hours in a 4.5 Liter Freeze Dry System (FreeZone, Labconco, Kansas City, MO, U.S.A.). We used a mortar and pestle to homogenise dried samples to a fine powder. To account for a potential bias introduced by lipids, which are ^13^C depleted compared to pure protein (Hussey et al., 2012; Sweeting et al., 2006), homogenised muscle samples were placed in 2 ml cryovials and suspended in a 2:1 chloroform-methanol mixture (CHCl_3_-CH_3_OH) for 24 hours at 30° C. Samples were centrifuged for 4 – 6 minutes, and the supernatant was removed. The process was repeated a second time. Samples were then left to dry with the cap of the cryovials open until all chloroform-methanol mixture was completely evaporated (after 24 – 48 hours). Sharks and rays accumulate urea and trimethylamine n-oxide (TMAO) in their tissues for osmoregulation (Ballantyne, 1997). These nitrogenous compounds, hereafter “urea”, are depleted in ^15^N and can therefore bias δ^15^N values. We removed urea following Li et al. (2016). In brief, homogenised samples were suspended in 1.9 ml deionised water, vortexed and left to rest for 24 hours before the samples were centrifuged, and the supernatant was removed. These steps were repeated three times before the samples were freeze-dried for a final time. Lipid- and urea-extracted samples at the University of Windsor were re-homogenised and between 0.1 and 0.6 mg of sample were weighed into tin capsules (5 x 8 mm) and analysed using a continuous-flow isotope ratio mass spectrometer (IMRS, Finnigan MAT Delta^Plus^, Thermo Finnigan, San Jose, California, U.S.A.). The mass spectrometer was coupled to an elemental analyser (Costech, Valencia, California, U.S.A.). The δ^13^C and δ^15^N values were normalised to the VPDB (Vienna Pee Dee Belemnite) and N^2^ (air), respectively.

Muscle and serum samples analysed at the University of Basel were first frozen at -80° C for 24 hours and then freeze-dried for 48 hours in a 4.5 Liter Freeze Dry System (FreeZone, Labconco, Kansas City, MO, U.S.A.). Dried samples were then homogenised to a fine powder using a mortar and pestle. Following Shipley et al. (2017), no lipid extraction was performed given expected low lipid content in serum. However, to account for differences in lipid extraction protocols, we compared the δ^13^C and δ^15^N values between lipid-and-urea-extracted and urea-extracted-only muscle samples and adjusted the values accordingly (see Supplementary Material, Fig S1, S2 A – B, and Table S1). Urea in muscle and serum samples analysed in Basel was removed following the steps described above. Urea extracted samples in Basel were re-dried in a drying oven at 60° C for 24 hours and re-homogenised. Between 0.4 and 0.6 mg of homogenised sample were weighted into tin capsules (5 x 9 mm, Säntis Analytical, Teufen, Switzerland) and analysed on a Flash 2000 elemental analyser coupled to a Thermo Finnigan Delta^Plus^ isotope ratio mass spectrometer (as above) via a Conflo IV interface (Thermo Fisher Scientific, Bremen, Germany). A thermal conductivity detector coupled to the elemental analyser was used to measure the amount of C and N (mg) in the sample prior to the introduction into the mass spectrometer, and the C:N ratio was calculated (all C:N ratios are reported in Tables S1 and S2). The δ^13^C and δ^15^N values were normalized to the VPDB and N^2^ scales, using laboratory standards (urea and sucrose powder for δ^13^C, caffeine powder and spirulina for δ^15^N) calibrated against international reference standards (IAEA-C3, USGS 40, IAEA-600, USGS 24, and IAEA-No-3).

Here we report the isotopic composition of muscle and serum samples using the standard delta (δ) notation (i.e. the relative difference in parts per mill [‰]) between the analysed sample and the used reference standards. The average analytical accuracy for samples analysed in Basel was ± 0.05 ‰ and ± 0.06 ‰ for δ^13^C and δ^15^N, respectively. Analytical precision, as determined by the analysis of duplicates (one duplicate per 10 samples): ± 0.2 ‰ δ^13^C and ± 0.1 ‰ δ^15^N for muscle samples analysed at the University of Windsor, ± 0.03 ‰ δ^13^C and ± 0.04 ‰ δ^15^N for muscle samples analysed at the University of Basel, and ± 0.04 ‰ δ^13^C and ± 0.04 ‰ δ^15^N for serum samples.

### 2.4. Condition metrics analysis

Serum samples used for the measurement of β-hydroxybutyrate, glucose and lactate concentrations, as well as serum osmolality, were analysed at the Department of Animal Health and Science Operations, at Disney’s Animal, Science & Environment in Lake Buena Vista, Florida, in the United States of America.

Frozen serum samples were thawed at room temperature prior to the analysis. Serum concentrations of β-hydroxybutyrate, hereafter “butyrates”, were measured in mmol/L using 10 μL of sample analysed in a STAT-Site® M analyser (EKF diagnostics, Boerne, Texas, U.S.A.). One butyrate measurement was discarded due to a NA reading. Glucose concentrations were measured in mg/dl by analysing 5 μL of serum in an Accu-Check® unit (Roche Diabetes Care, Inc., Indianapolis, Indiana, U.S.A). Lactate concentrations were measured in mmol/L by analysing 5 μL of serum in a Lactate Scout® unit (EKF diagnostics, Boerne, Texas, U.S.A.). Serum osmolality was measured in mOsm using 5 μL of serum in an Osmette® III Osmometer (Precisions Systems, Natick, Massachusetts, U.S.A.).

Our interpretation of condition metrics for wild caught individuals was based on preliminary findings of blood biochemical analyses in aquarium-managed southern stingrays (Mylniczenko et al., unpublished data). Butyrate concentrations ≤ 0.7 mmol/L were interpreted as normal; Butyrate concentrations > 0.7 mmol/L indicate that the animal was mobilising energy. Glucose concentrations between 10 – 40 mg/dl were viewed as normal, and an upper threshold of 60 mg/dl was used, with values above this threshold indicating an active stress response and mobilisation of hepatic glycogen storages. Lactate concentrations of < 3 mmol/L were interpreted as normal, while concentrations between 3 – 10 mmol/L signalled muscle exertion and hypoventilation. Animals with lactate concentrations > 12 mmol/L were expected to be physiologically compromised. Blood osmolality values of > 900 mOsm (≈ 32 ppt) and > 1000 mOsm (≈ 35 ppt) correspond to the iso- and hyperosmotic range compared to seawater. Osmolality values < 900 mOsm were used as an indication of an animal experiencing low urea levels which we interpret as representative of low lipids and consequently a smaller liver (i.e. reduced condition).

### 2.5. Statistical analysis

Prior to statistical comparisons of δ^13^C and δ^15^N values, we accounted for fractionation (i.e. the difference in isotopic composition between southern stingrays and their prey) by applying diet-tissue discrimination factors (DTDFs; Hobson & Clark, 1992; Hussey, MacNeil, et al., 2012). We applied DTDFs documented in captive leopard sharks, *Triakis semifasciata*, fed with squid, *Loligo opalescencs* (Kim et al., 2012). For muscle samples we used a DTDF of 1.7 ‰ for Δ^13^C, and 3.7 ‰ for Δ^15^N. For serum we used a DTDF of 2.8 ‰ for Δ^13^C and 2.2 ‰ for Δ^15^N. Raw isotopic data are reported in the Supplementary Material (Table S1 and S2). The δ^13^C and δ^15^N values of southern stingrays younger than 1 year are reported in the Supplementary Material but were excluded from statistical analyses as they are influenced by maternal contributions and do not reflect their own diet (Olin et al., 2011).

To examine variation in carbon and nitrogen isotope data and condition metrics among individuals, we investigated the effects of sex, capture location, season and size by simulating 5000 parameter values from the joint posterior distribution of linear models (LMs) and linear mixed models (LMMs), which were formulated using the *lme4* R package (version 1.1-31; Bates et al., 2015). Based on water temperature, we defined two captures seasons: (1) summer (i.e. June – November) and (2) winter (i.e. December – May). Season terminology was chosen to maintain consistency and allow for comparability with prior work in this study system (van Zinnicq Bergmann et al., 2022). We used the capture location as a proxy for predator exposure. Based on tissue-specific metabolic turnover rates of δ^13^C and δ^15^N and different underlying physiological processes of the condition metrics, we formulated different statistical models for isotopic ratios in muscle and serum samples, and for the condition metrics butyrates, glucose, lactate and osmolality (see Table 1). Given the shorter isotopic turnover rates in blood (MacNeil et al. 2006), we included the categorical variable season in an interaction with location in the LMMs assessing patterns in statistical models for serum samples only. The values of the joint posterior distribution were simulated using the *arm* R package (version 1.13-1; Gelman & Su, 2022). We used the simulated values to calculate the 2.5 % and 97.5 % quantile of the marginal posterior distribution of our model parameters and based statistical inference on the resulting 95 % CrI.

**Table 1.** Table overview of variables and formulas used in the statistical models exploring different effects on documented stable isotope ratios and condition metrics. Linear Mixed Models (LMMs) exploring δ^13^C and δ^15^N values from muscle samples included the fixed effects sex, location and disc width. To control for individual variation, we used the unique PIT tag number of sampled rays as random effect in all models, except for the butyrate, glucose and lactate models, where the variance of the random effect was negligible, and a LM without the random effect was formulated to improve the model fit. The abbreviations in the explaining variable location stand for: “LAG” – lagoon, “BFH” – Bonefish Hole, and “SB” – South Bimini.

**A: Variables**
| *Name* | *Description* | *Unit* |
| --- | --- | --- |
| $\delta^{13}\text{C}$ | Ratio of $^{13}\text{C}$ to $^{12}\text{C}$ in muscle and serum samples relative to international reference standards | ‰ |
| $\delta^{15}\text{N}$ | Ratio of $^{15}\text{N}$ to $^{14}\text{N}$ in muscle and serum samples relative to international reference standards | ‰ |
| Butyrate | Butyrate concentration in serum samples | mmol/L |
| DW | Disc width of the sampled southern stingrays | cm |
| Glucose | Glucose concentration in serum samples | mg/dl |
| Lactate | Lactate concentration in serum samples | mmol/L |
| Location | Capture location of southern stingrays (see Fig. 1) | Categorical, 3 levels, “BFH”, “LAG” and “SB” |
| Osmolality | Osmolality of serum samples | mOsm |
| PIT | Unique passive integrated transponder (PIT) tag number of sampled southern stingrays | n/a |
| Season | Season of capture/sampling of southern stingrays | Categorical, 2 levels, “summer” and “winter” |
| Sex | Sex (female or male) of sampled southern stingrays | Categorical, 2 levels, “F” and “M” |

Bayesian isotope mixing models were implemented via the *MixSIAR* R package (version 3.1.12; Stock et al., 2018) to explore the relative contribution of different prey groups to the diet of southern stingrays. We built an isotopic mixing space (i.e. isospace) by calculating the mean δ^13^C and δ^15^N values ± SD across potential prey species (Table S3) spanning four main prey groups: annelids (δ^13^C: -15.8 ± 7.8 ‰, δ^15^N: 5.3 ± 2.2 ‰), crustaceans (δ^13^C: -16.2 ± 3.1 ‰, δ^15^N: 6.5 ± 2.4 ‰), molluscs (δ^13^C: -13.5 ± 2.5 ‰, δ^15^N: 4.1 ± 0.7 ‰), and teleosts (δ^13^C: -18.7 ± 5.5 ‰, δ^15^N: 7.9 ± 2.6 ‰). Prey groups were chosen based on the available literature discussing stomach contents and diet preferences of southern stingrays (Bigelow & Schroeder, 1953; Bowman et al., 2000; Gilliam & Sullivan, 1993; O’Shea et al., 2020; Randall, 1967; Smith & Herrnkind, 1992; Snelson Jr. & Williams, 1981; Tilley et al., 2013). Because of the slow isotopic turnover rate of elasmobranch muscle tissue (MacNeil et al., 2006), we only included δ^13^C and δ^15^N values from serum samples in the mixing models. We fitted separate models for mature and immature female individuals during the summer and winter, and mature males during summer and winter. Each model consisted of three chains, and we chose chain lengths of 1 000 000 iterations with a thinning interval of 500 and we discarded the first 500 000 iterations. Model convergence was assessed using Gelman-Rubin diagnostics (Stock et al., 2018) and visual inspection of the trace and density plots.

Lastly, Bayesian correlation tests were performed between the δ^13^C and δ^15^N values and the four condition metrics in serum samples using the *bayestestR* R package (version 0.17.0; Makowski et al. 2019) to explore potential relationships between the trophic ecology and the condition of sampled individuals. We sampled 5000 iterations from the posterior distribution to calculate the 2.5 % and 97.5 % quantiles and based statistical inference on the resulting 95 % CrI In case of the the 95 % CrI not including 0, we used the Bayes Factor as relative evidence in favor of a possible correlation based on Jeffreys (1961), where BF = 1 is no, BF > 1 and <=3 is anecdotal, BF > 3 and <= 10 is moderate, BF > 10 and <= 30 is strong, BF > 30 and <= 100 is very strong, and BF > 100 is extreme evidence in favour of a correlation between two variables.

All statistical analyses were performed in R (version 4.2.2; R Core Team, 2022) and summarised data are reported as mean ± standard deviation (SD).

### 2.6. Ethical note

This study and all described research activities, capture and sampling protocols were conducted and approved under permits MAMR/LIA/22 and MA&MR/FIS/17B issued by the Department of Marine Resources (DMR) and the Department of Environmental Planning and Protection (DEPP) of The Bahamas.

Following capture and during the sampling protocol, southern stingrays were constantly monitored, and all rays were released alive and competent for release. Following release, all southern stingrays were followed by boat or by walking to ensure a good release condition. If an individual settled on the sediment to rest, we monitored its buccal pumping activity and we stayed with the animal until it resumed swimming on its own.

## 3. Results

Between July 10^th^, 2018, and November 27^th^, 2019, we sampled 165 unique southern stingrays (147 females and 18 males) across 197 captures (131 captures during summer, 66 captures during winter, Tables S1, S2 and S4). Eleven individuals were sampled more than once for muscle (range: 2 – 3 samples per individual, Table S1) and serum (range: 2 – 3 samples per individual, Table S2) resulting in 152 muscle (123 females and 17 males) and 144 serum samples (119 females and 12 males). The disc width of sampled females ranged from 39.6 to 108.4 cm (mean ± SD: 82.3 ± 14.4 cm) and from 42.6 to 68.2 cm (mean ± SD: 50.6 ± 6.0 cm) in sampled males. Based on published disc-width-at-age values (Hayne et al., 2018; Henningsen & Leaf, 2010), 112 females were sexually mature, 33 were sexually immature, and two individuals were characterised as young-of-the-year and thus removed from further analyses. All male individuals were sexually mature, but no male individuals were captured in Bonefish Hole. (Table S1, S2 and S4).

### 3.1. Stable isotope analysis

DTDF-corrected muscle isotope values ranged from -16.8 to -10.5 ‰ (mean ± SD: -13.1 ± 1.3 ‰, N = 149) for δ^13^C and from 2.2 to 7.5 ‰ for δ^15^N (mean ± SD: 4.0 ± 0.7 ‰, N = 149, Fig. 2 A, see Table S1 for raw isotopic ratios). The differences in muscle isotope values from re-captured and re-sampled individuals ranged from 0.1 – 1.8 ‰ (mean ± SD: 0.5 ± 0.6 ‰, N = 12) for δ^13^C and from 0.1 – 1.5 ‰ (mean ± SD: 0.6 ± 0.4 ‰, N = 12) for δ^15^N (Table S1). Muscle δ^13^C values were not different between females (mean ± SD: -13.2 ± 1.3 ‰, N = 132) and males (mean ± SD: -12.4 ± 1.2 ‰, N = 17, Fig. S3 A, Table 2), nor between the capture locations (Fig. 3 A – C, Table 2). We found that δ^13^C values were, albeit very weekly (posterior mean: -0.018, 95 % CrI: [-0.035, -0.002], Table 2), negatively correlated with disc width (Fig. 3 A – C). Muscle δ^15^N values in females (mean ± SD: 3.9 ± 0.7 ‰, N = 132) and males (mean ± SD: 4.45 ± 0.8 ‰, N = 17) were not significantly different (Fig. S3 B, Table 2), but muscle δ^15^N values were higher for females sampled in the lagoon and off South Bimini compared to females sampled in Bonefish Hole (Fig. 3 D – F, Fig S3 B, Table 2). In contrast to δ^13^C values, the disc width of southern stingrays had no significant effect on the δ^15^N values in muscle samples (Fig. 3 D – E, Table 2).

**Fig. 2.**
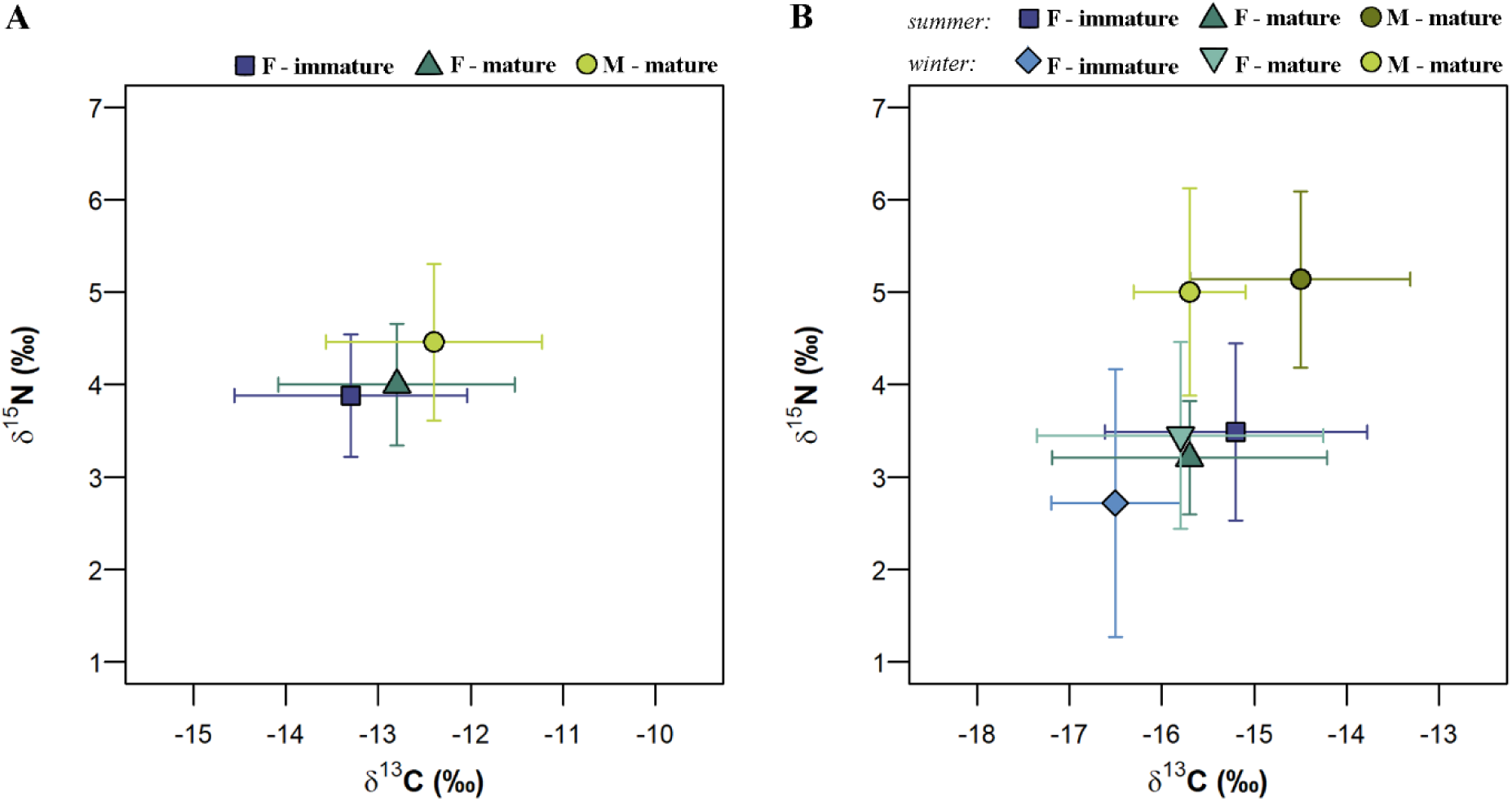
Mean δ^13^C and δ^15^N values corrected for diet-tissue discrimination factors (DTDF) for (A) white muscle tissue and (B) serum samples. Isotopic values of females are shown separately for immature (squares) and mature (triangles) individuals. All sampled males (circles) were determined to be mature based on calcified claspers. Isotopic values from serum samples in panel B are shown for the two seasons summer (darker colours) and winter (lighter colours). Symbols show the mean values, and the error bars show ± 1 standard deviation (SD).

**Fig. 3.**
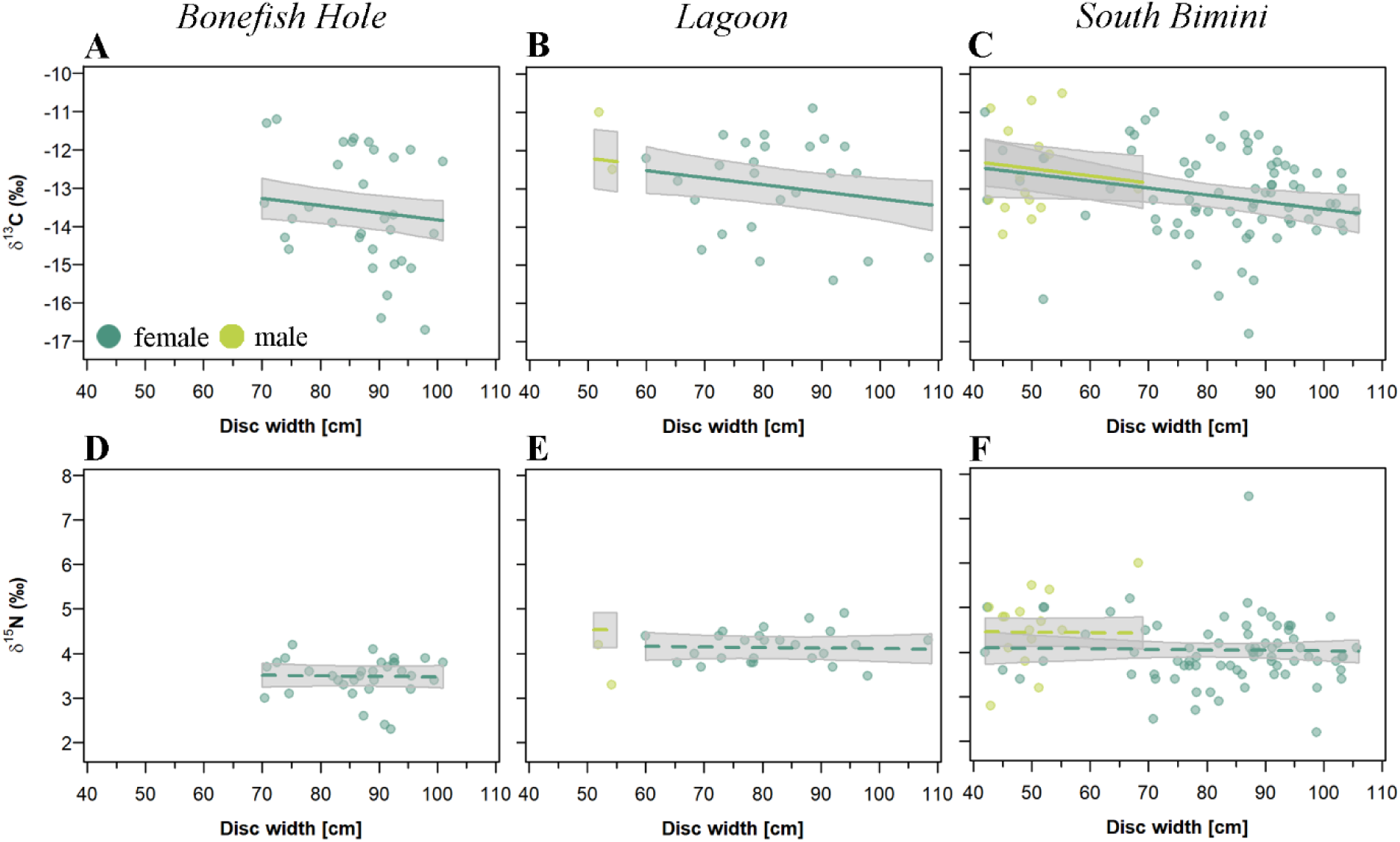
Effects of sex and disc width on (A – C) δ^13^C and (D – F) δ^15^N values from muscle samples collected in Bonefish Hole, the lagoon, and South Bimini. Coloured points show the raw isotopic ratios by sex. Coloured lines show the regression coefficient of disc width on the δ^13^C and δ^15^N values in females and males as extracted from the corresponding LMM. Southern stingrays display a sexual dimorphism with females growing to substantially larger sizes (maximum disc width: ∼ 150 cm) than males (maximum disc width: ∼ 67 cm; Henningsen & Leaf, 2010; Last et al., 2016). Solid lines show a clear effect, dashed lines show trends, or no effects based on the 95 % CrIs of the posterior, which are shown as grey polygons.

**Table 2.**
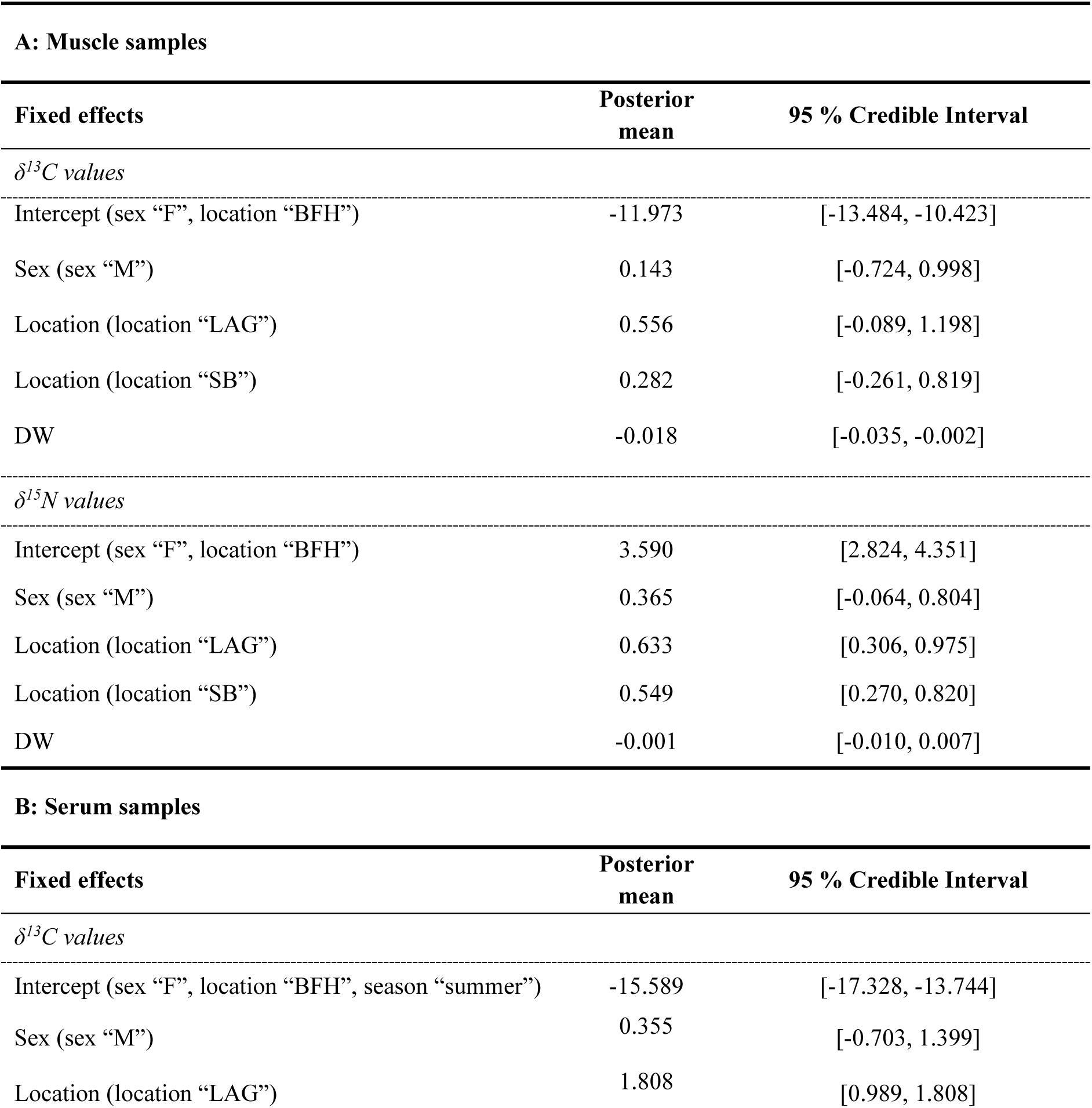
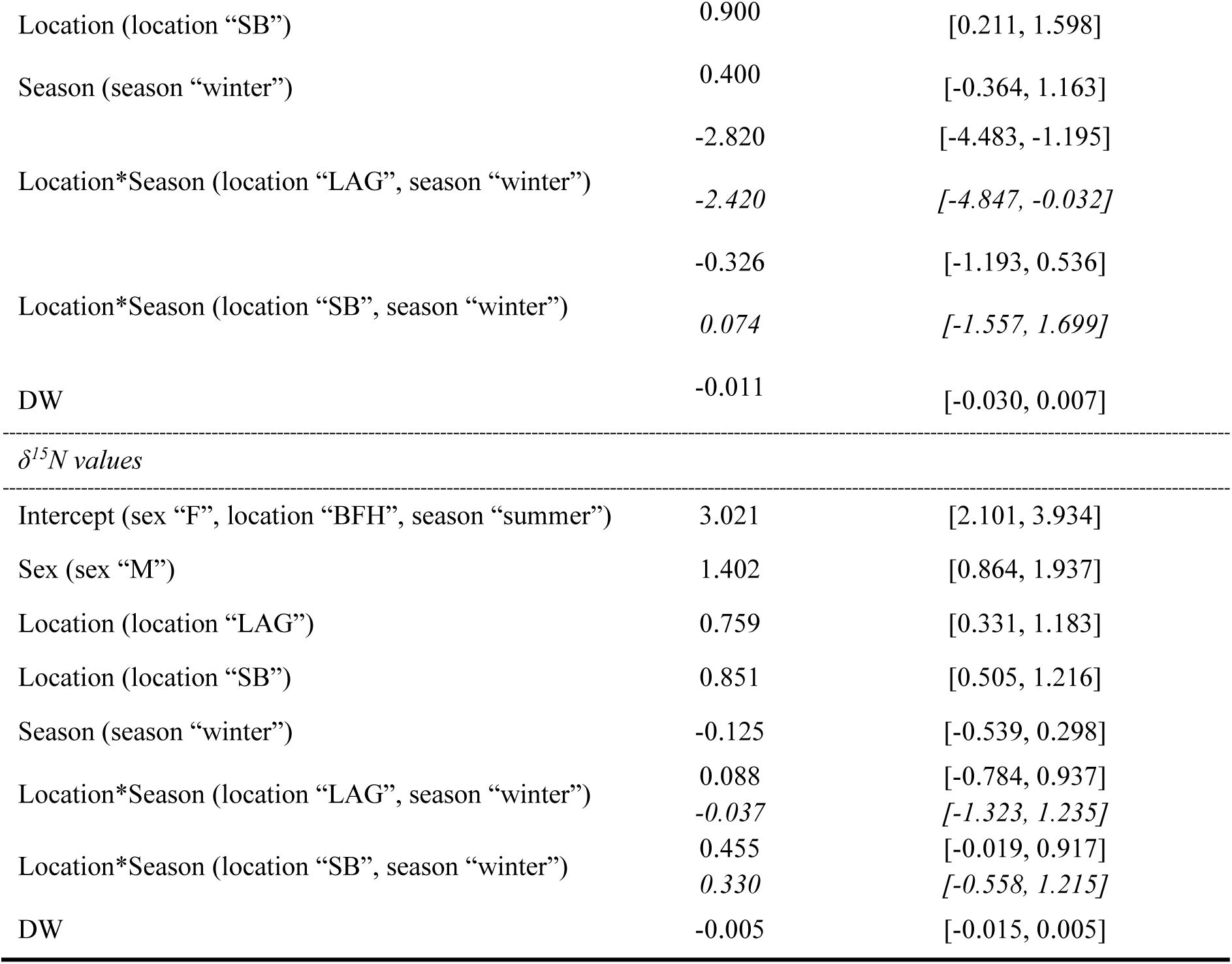
Posterior means and 95 % credible intervals (CrIs) of fixed effects in linear mixed models (LMMs) exploring δ^13^C and δ^15^N values in (A) muscle and (B) serum samples. Calculated coefficients in interactions are shown in italic. Factor levels of coefficients in the “Fixed effects” column are shown in parentheses. DW stands for disc width. Values were rounded three digits after the decimal point. The 95 % CrIs are written in brackets with the first element showing the 2.5 %, and the second element showing the 97.5 % quantile. The abbreviations in the explaining variable location stand for: “LAG” – lagoon, “BFH” – Bonefish Hole, and “SB” – South Bimini.

The DTDF-corrected serum δ^13^C and δ^15^N values ranged from -19.8 to -12.5 ‰ (mean ± SD: -15.6 ± 1.5 ‰, N = 142) and from 1.4 to 7.2 ‰ (mean ± SD: 3.5 ± 1.0 ‰, N = 142, Fig. 2 B, see Table S2 for raw isotopic ratios), respectively. The differences in serum δ^13^C values from recaptured individuals that were resampled during the same season ranged from 0.0 – 0.8 ‰ (mean ± SD: 0.3 ± 0.4 ‰, N = 3) during summer and from 0.3 – 0.6 ‰ (mean ± SD: 0.4 ± 0.2 ‰, N = 3) during winter (Table S2). The serum δ^15^N values from recaptured individuals that were resampled during the same season ranged from 0.0 – 0.3 ‰ (mean ± SD: 0.1 ± 0.2 ‰, N = 3) during summer and from 0.2 – 0.4 ‰ (mean ± SD: 0.3 ± 0.1 ‰, N = 3) during winter (Table S2). There was no difference between serum δ^13^C values of females and males (Fig. S3 C, Table 2). We found no seasonal differences in serum δ^13^C samples collected in Bonefish Hole and off South Bimini, but in the lagoon, we found lower δ ^13^C values during the winter compared to summer (Fig. 4 A – C, Table 2). As a result, serum δ^13^C values of southern stingrays caught off South Bimini (Fig. 4 C) were the same as those for individuals sampled in the lagoon during the summer (Fig. 4 B), but were higher when compared to δ^13^C values for animals sampled from the lagoon during winter and from Bonefish Hole during both seasons (Fig. 4 A and B, Table 2). In contrast to muscle tissue data, there was, however, no trend towards lower serum δ^13^C values with increasing disc width (Fig. 4 A – C, Table 2).

**Fig. 4.**
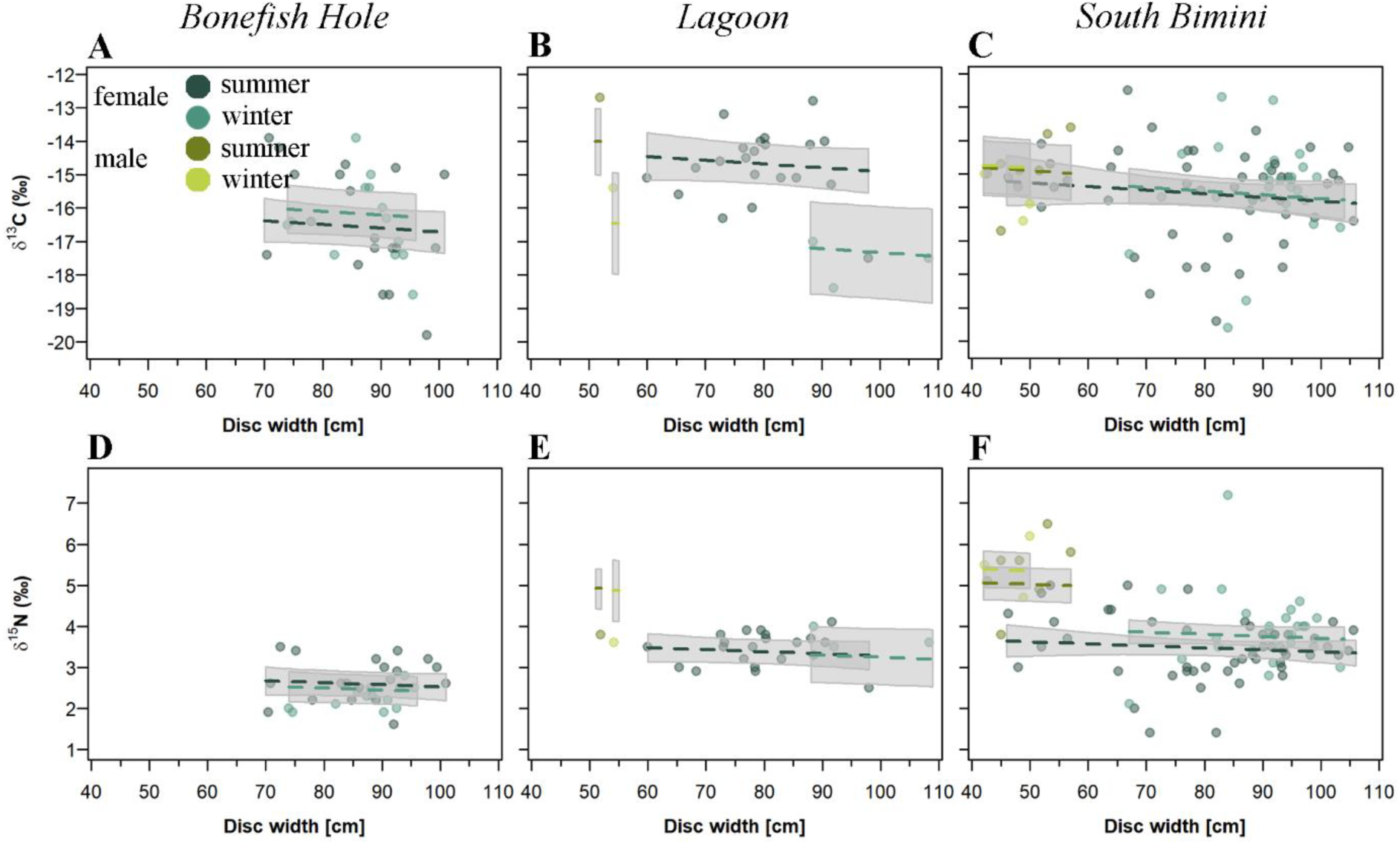
Effects of sex, season, and disc width on (A – C) δ^13^C and (E – F) δ^15^N values from serum samples collected in Bonefish Hole, the lagoon, and South Bimini. Coloured lines show the regression coefficient of disc width on the δ^13^C and δ^15^N values in females and males during summer (darker colours) and winter (brighter colours) as extracted from the corresponding LMM. Solid lines show a clear effect, dashed lines show trends, or no effects based on the 95 % CrIs of the posterior, which are shown as a grey polygons.

Serum δ^15^N values were lower in females compared to males (Fig. S3 D, Table 2). There was no seasonal effect on serum δ^15^N values at the three sampling locations, but we found higher serum δ ^15^N values in the lagoon and South Bimini compared to those from Bonefish Hole (Fig. 4 D – E, Table 2). There was no effect of size on serum δ^15^N values (Fig. 4 D – E, Table 2).

Generally, molluscs represented the dominant prey group for southern stingrays followed by annelids (Fig. 5 A - F), except for mature females, where molluscs and annelids were equally common across summer and winter (Fig. 5 C, D). Crustaceans and teleosts contributed less towards the diet of southern stingrays. However, in immature females we found a shift in the estimated diet between the two seasons (Fig. 5 A, B): molluscs remained the dominant prey group across both seasons, but crustaceans and teleosts became more common during winter, with a decrease in the importance of annelids. The estimated diet of mature males (Fig. 5 E, F) was similar to the diet of immature females during winter – molluscs were the dominant prey group, and crustaceans, teleosts and annelids were less common. Equivalent to mature females, the estimated diet patterns of males were stable between seasons.

**Fig. 5.**
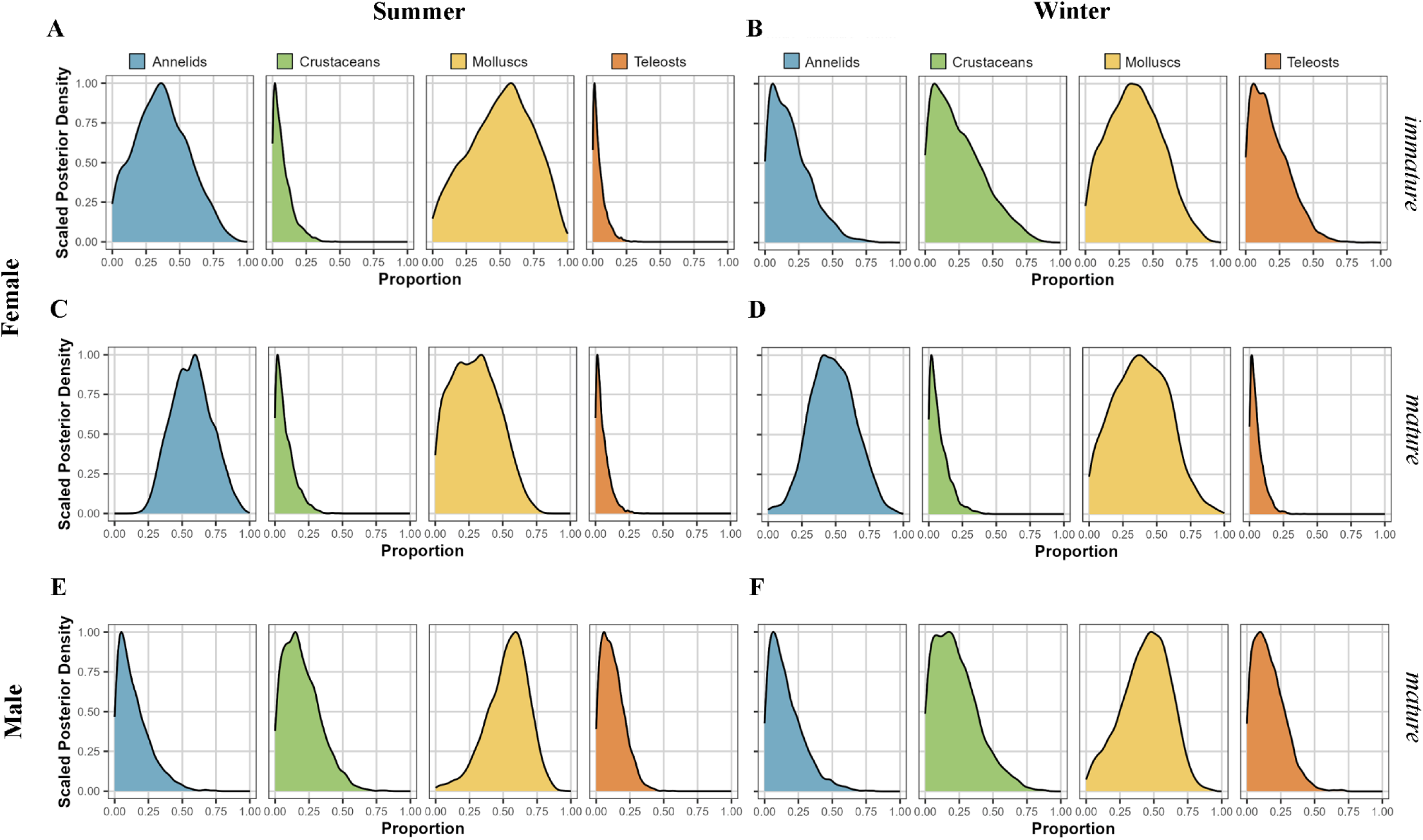
Scaled posterior densities extracted from the Bayesian mixing models estimating the relative contribution of different prey groups to the diet of southern stingrays. The scaled posterior density of the proportion of annelids, crustaceans, molluscs and teleosts is shown for immature female stingrays during the (A) summer and (B) winter, mature female stingrays during the (C) summer and (D) winter, and mature male stingrays during the (E) summer and (F) winter.

### 3.2. Condition metrics

We analysed 122 serum samples from 114 unique individuals (112 females, 10 males) to determine serum butyrate, lactate and glucose concentrations and osmolality (Table S4). The disc width of sampled females and males ranged from 52.0 – 108.4 cm (mean ± SD: 84.9 ± 11.6 cm) and from 42.2 to 54.2 cm (mean ± SD: 48.4 ± 4.41 cm), respectively (Table S4).

Measured butyrate concentrations were 0.56 ± 0.34 mmol/L for females (range 0.40 – 2.50 mmol/L) and 0.56 ± 0.16 mmol/L for males (range: 0.40 – 0.80 mmol/L) and were similar between sexes, across seasons and locations (Fig. 6 A - C, Fig. S4 A, Table 3). Butyrate concentrations did not change with increasing disc width of the sampled individuals (Fig. 6 A - C, Table 3). In six samples we measured butyrate concentrations above 0.7 mmol/L (4 females, 2 males, Table S4).

**Fig. 6.**
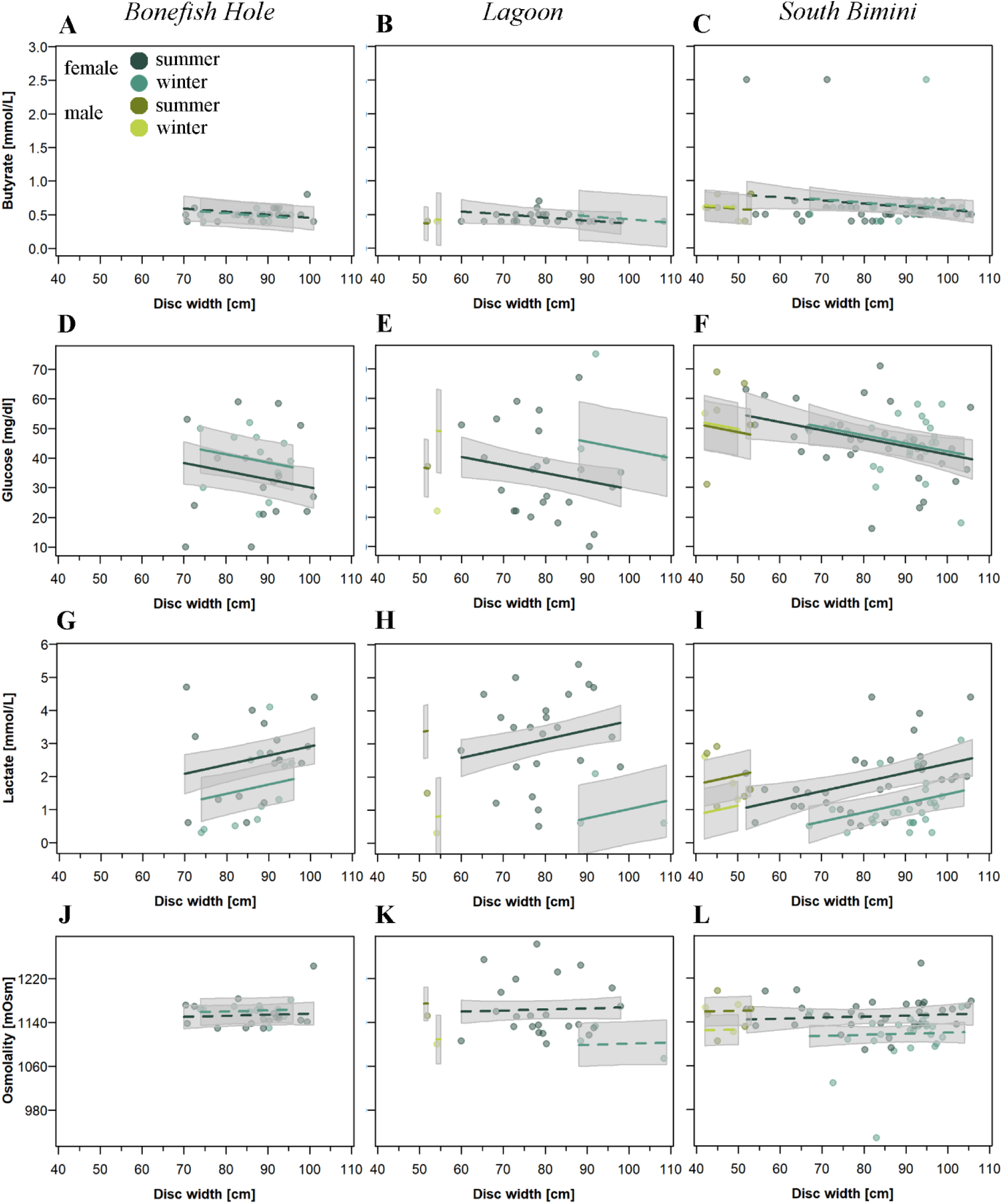
Effects of sex, season, and disc width on concentrations of (A – C) butyrates, (D – F) glucose, (G – I) lactate, and (J – L) osmolality of serum samples collected in Bonefish Hole, the lagoon, and South Bimini. Coloured lines show the regression coefficient of disc width on the body condition metrics of females and males during summer (darker colours) and winter (brighter colours) as extracted from the corresponding statistical model. Solid lines show a clear effect, dashed lines show trends, or no effects based on the 95 % CrIs of the posterior, which are shown as a grey polygons.

**Table 3.** Posterior means and 95 % credible intervals (CrIs) of fixed effects in statistical models exploring butyrate, glucose and lactate concentrations in, and osmolality of, serum samples. Calculated coefficients in interactions are shown in italic. Factor levels of coefficients in the “Fixed effects” column are shown in parentheses. DW stands for disc width. Values were rounded three digits after the decimal point. The 95 % CrIs are written in brackets with the first element showing the 2.5 %, and the second element showing the 97.5 % quantile. The abbreviations in the explaining variable location stand for: “LAG” – lagoon, “BFH” – Bonefish Hole, and “SB” – South Bimini.

| Condition metrics |  |  |
| --- | --- | --- |
| Fixed effects | Posterior mean | 95 % Credible Interval |
| *Butyrate [mmol/L]* |  |  |
| Intercept (sex “F”, location “BFH”, season “summer”) | 0.913 | [0.411, 1.416] |
| Sex (sex “M”) | -0.215 | [-0.519, 0.080] |
| Location (location “LAG”) | -0.093 | [-0.299, 0.116] |
| Location (location “SB”) | 0.114 | [-0.072, 0.304] |
| Season (season “winter”) | -0.024 | [-0.273, 0.224] |
| Location*Season (location “LAG”, season “winter”) | 0.091<br>0.067 | [-0.384, 0.562]<br>[-0.657, 0.786] |
| Location*Season (location “SB”, season “winter”) | 0.038<br>0.014 | [-0.254, 0.343]<br>[-0.527, 0.567] |
| DW | -0.005 | [-0.010, 0.001] |
| *Glucose [mg/dl]* |  |  |
| Intercept (sex “F”, location “BFH”, season “summer”) | 57.626 | [37.549, 77.779] |
| Sex (sex “M”) | -6.156 | [-17.784, 5.632] |
| Location (location “LAG”) | -0.866 | [-8.957, 6.848] |
| Location (location “SB”) | 10.951 | [3.498, 18.325] |
| Season (season “winter”) | 5.637 | [-3.788, 15.168] |
| Location*Season (location “LAG”, season “winter”) | 7.735<br>13.372 | [-9.364, 24.882]<br>[-13.152, 40.050] |
| Location*Season (location “SB”, season “winter”) | -4.616<br>1.021 | [-15.929, 7.073]<br>[-19.717, 22.241] |
| DW | -0.275 | [-0.497, -0.052] |
| *Lactate [mmol/L]* |  |  |
| Intercept (sex “F”, location “BFH”, season “summer”) | 0.139 | [-1.587, 1.768] |
| Sex (sex “M”) | 1.037 | [0.084, 2.004] |
| Location (location “LAG”) | 0.780 | [0.128, 1.436] |
| Location (location “SB”) | -0.529 | [-1.137, 0.065] |
| Season (season “winter”) | -0.876 | [-1.682, -0.876] |
| Location*Season (location “LAG”, season “winter”) | -1.789<br>-2.665 | [-3.236, -0.362]<br>[-4.918, -0.435] |
| Location*Season (location “SB”, season “winter”) | -0.040<br>-0.916 | [-0.993, 0.909]<br>[-2.675, 0.836] |
| DW | 0.028 | [0.010, 0.046] |

| *Osmolality [mOsm]* |  |  |
| --- | --- | --- |
| Intercept (sex “F”, location “BFH”, season “summer”) | 1136.392 | [1075.077, 1198.491] |
| Sex (sex “M”) | 16.324 | [-20.242, 52.259] |
| Location (location “LAG”) | 11.256 | [-13.358, 36.683] |
| Location (location “SB”) | -2.240 | [-24.779, 20.467] |
| Season (season “winter”) | 8.337 | [-20.613, 38.141] |
| Location*Season (location “LAG”, season “winter”) | -74.353 | [-126.224, -21.864] |
|  | -66.016 | [-146.837, 16.277] |
| Location*Season (location “SB”, season “winter”) | -42.101 | [-75.841, -8.348] |
|  | -33.764 | [-96.454, 29.793] |
| DW | 0.196 | [-0.499, 0.867] |

Serum glucose concentrations ranged from 10.00 to 75.00 mg/dl in females (mean ± SD: 40.90 ± 13.57 mg/dl, N = 112) and from 22.00 – 69.00 mg/dl in males (mean ± SD: 48.40 ± 14.68 mg/dl, N = 10) but there was no difference between the two sexes (Fig. 6 A – E, Fig. S4 B, Table 3). Glucose concentrations were similar during summer and winter, for animals collected in Bonefish Hole and the lagoon (Fig. 6 D and E, Table 3) and for those collected in the lagoon and off South Bimini (Fig. 6 E and F, Table 3). Glucose concentrations were, however, higher for animals collected off South Bimini compared to those in Bonefish Hole (Fig. 6 D and F, Table 3). We found a decrease in glucose concentrations with increasing disc widths (Fig. 6 D – F, Table 3). Serum glucose concentrations > 60 mg/dl indicating an acute stress response were recorded in eight individuals (6 females, 2 males, Table S4).

Lactate concentrations of serum for female southern stingrays was 2.04 ± 1.31 mmol/L (range: 0.30 – 5.40 mmol/L, N = 112) and for males was 1.72 ± 0.81 mmol/L (range: 0.3 – 2.9 mmol/L, N = 10, Table S4). Measured lactate concentrations were highest in samples from the lagoon. Concentrations were generally lower for samples collected during the winter, except for South Bimini (Fig. 6 G – I, Fig. S4 C, Table 3). Serum lactate concentrations increased with increasing disc width and were higher in males (Fig. 6 G – I, Table 3). Serum lactate concentrations in 26 samples indicated hypoventilation and muscle exertion (3.00 – 10.00 mmol/L), but there was no indication of individuals experiencing a physiological crisis (> 12 mmol/L, Table S4).

We found no difference for serum osmolality between female (mean ± SD: 1146.53 ± 42.78. mOsm, range: 928 – 1283 mOsm, N = 112) and male southern stingrays (mean ± SD: 1146.10 ± 30.94 mOsm, range: 1100 – 1197 mOsm, N = 10, Fig. S4 D, Table 3). Serum osmolality was similar between samples collected during summer and winter, and between all three locations (Fig. 6 J – K, Table 3). However, there appeared to be a trend towards lower osmolality in serum samples during the winter compared to summer in individuals from the lagoon (Fig. 6 K). There was no effect of disc width on measured serum osmolality (Fig. 6 J – K, Table 3).

### 3.3. Correlation between stable isotope ratios and condition metrics

There was no correlation between serum δ^13^C values and any of the condition metrics (Table 4). Similarly, there was no correlation between serum δ^15^N values, butyrate, and lactate concentrations. We found moderate evidence (BF = 9.29) for a weak positive correlation between serum δ^15^N values and glucose, and anecdotal evidence (BF = 2.27) for a weak negative correlation between serum δ^15^N values and serum osmolality (Table 4).

**Table 4.** Bayesian correlation tests between stable isotope ratios and condition metrics. Values were rounded three digits after the decimal point. The 95 % Credible Intervals (CrIs) are written in brackets with the first element showing the 2.5, and the second element showing the 97.5 % quantile. The Bayes Factor (BF) is interpreted based on Jeffreys (1961) as relative evidence in favor (BF > 1).

| Isotope | Condition metric | Posterior mean | BF | Relative evidence |
| --- | --- | --- | --- | --- |
| $\delta^{13}\text{C}$ | Butyrate | 0.065<br>[-0.101, 0.240] | 0.273 | NA |
|  | Glucose | 0.084<br>[-0.081, 0.255] | 0.328 | NA |
|  | Lactate | -0.095<br>[-0.264, 0.085] | 0.367 | NA |
|  | Osmolality | -0.051<br>[-0.230, 0.123] | 0.249 | NA |
| $\delta^{15}\text{N}$ | Butyrate | 0.166<br>[-0.013, 0.355] | 1.295 | NA |
|  | Glucose | 0.240<br>[0.079, 0.412] | 9.292 | Moderate evidence in favor |
|  | Lactate | -0.073<br>[-0.250, 0.095] | 0.293 | NA |
|  | Osmolality | -0.190<br>[-0.347, -0.014] | 2.271 | Anecdotal evidence in favor |

## 4. Discussion

Here, the analysis of carbon (δ^13^C) and nitrogen (δ^15^N) stable isotopes in white muscle tissue and serum samples of southern stingrays revealed relatively uniform trophic interactions across seasons, with the exception of lower muscle δ^13^C values for larger individuals, and higher serum δ^15^N values for males. Bayesian mixing models identified a dietary preference for molluscs, but a relatively wide range of δ^13^C and δ^15^N values were recorded for both sexes, as is typical for mesopredators. Our results underscore the southern stingrays’ role as an ecological link in the food web feeding on a variety of prey at multiple trophic levels (O’Gorman & Emmerson, 2009; Tilley et al., 2013). Further, we provide baseline data for four condition metrics in southern stingrays – serum concentrations of β-hydroxybutyrate, glucose, lactate and osmolality. Through this multi-tiered approach, we contribute towards a better understanding of the condition and functional roles of wild caught southern stingrays in marine ecosystems.

### 4.1. Trophic ecology

Overall, we documented a very high variability in δ^13^C values at the individual-level. However, all values in this study fell between the range of δ^13^C values typical for mangrove (more depleted in ^13^C) and seagrass (less depleted in ^13^C) habitats as previously reported for Bimini (Hussey et al., 2017). Except for southern stingrays sampled in the lagoon, we observed temporally stable δ^13^C values based on sampling seasons and tissues analysed, indicating southern stingrays used the same basal energy sources during summer and winter in Bonefish Hole and off South Bimini (Peterson & Fry, 1987). The seasonal differences and the slightly smaller variability in δ^13^C values in the lagoon may be explained by biogeochemical processes or environmentally impacted fractionation patterns know to alter isotopic signatures (Barnes et al., 2007; Freeman, 2001). However, the seasonal patterns could also be a function of varying seagrass coverage in the lagoon between summer and winter (van Tussenbroek et al., 2014), where southern stingrays predominantly forage in seagrass habitats during summer and switch to mangrove habitats during the winter when seagrass coverage is reduced. This selectivity between frequented habitats during the different seasons in the lagoon may be facilitated by an overall lower predation risk in this area, whereas the increased individual diet diversity as observed in Bonefish Hole and off South Bimini can be a consequence of higher predation risk, when prey limit their movements as an avoidance strategy, restricting their foraging opportunities (Catano et al., 2014; Reddin et al., 2016).

Here, documented δ^13^C values changed with southern stingray size, and decreased with increasing disc width, similar to southern stingrays in Exuma and Eleuthera (O’Shea et al., 2020). This ontogenetic shift towards lower δ^13^C in larger individuals indicates a shift towards more mangrove-tied prey and could be driven by pregnant individuals that seek out these shallow, sheltered and warm water habitats favourable for reproduction (Hight & Lowe, 2007; Jirik & Lowe, 2012; Tokunaga et al., 2022), as we frequently caught large, sexually mature and pregnant female southern stingrays near the mangroves. The shift towards lower δ^13^C values could also be due to larger southern stingrays relying on more molluscs in their diet, which can be strongly depleted in ^13^C due to chemosynthetic processes (Murchie et al., 2019). However, mixing model outputs in this study indicated that molluscs are important prey across all sizes, making this explanation unlikely. Additionally, the correlation between disc width and δ^13^C values was very weak and only observable in muscle samples. We therefore suggest interpreting the effect of disc width cautiously as the biological implications of these findings may be limited within our study system.

The δ^15^N values of southern stingrays in Bimini were comparable with values for other southern stingrays sampled in The Bahamas and Belize (O’Shea et al., 2020; Shipley et al., 2017, 2018, 2023; Tilley et al., 2013), but lower than in other mesopredatory elasmobranchs and large teleosts frequently observed in The Bahamas. This suggests mesopredators form a multi-tiered functional group (Bond et al., 2018; Shipley et al., 2018, 2023; Zhu et al., 2019). An ontogenetic shift towards the consumption of higher trophic level prey with increasing body size is frequently described for elasmobranchs and teleosts (Cocheret de la Morinière et al., 2003; Eggleston et al., 1998; Lucifora et al., 2009; Matich et al., 2019; Vaslet et al., 2011), but this was not observed among southern stingrays in Bimini based on δ^15^N values. Ontogenetic shifts towards higher δ^15^N values with increasing size often relate to size-based shifts in jaws and/or increased speed and agility that enable the capture of faster prey (McElroy et al., 2006; Papastamatiou et al., 2006; White et al., 2004). However, southern stingrays are anatomically designed to feed on prey living on or in the sediment, with our mixing models showing a preference for annelids and molluscs, which are likely not gape limiting in this species. Consequently, we would not expect an ontogenetic shift towards higher δ^15^N values. It is also possible that smaller southern stingrays eat smaller body sized molluscs, and a lack of dietary change in molluscs with size (herbivores vs. detritivores) would lead to a non-observable dietary change in the consumer.

We found, however, higher δ^15^N values in males, which on average were smaller than the caught females. Males and females feeding at different trophic levels could be a consequence of female and male southern stingrays partitioning resources in Bimini, thereby reducing intraspecific agonistic behaviour, including competition. Additionally, and supported by our capture data, males may spend less time in the shallow, protected waters, and may show increased movement and larger space use areas when looking for mating opportunities, thus inhabiting different habitats and foraging on prey with higher δ^15^N values.

Our estimated proportions of annelids, crustaceans, molluscs and teleosts in the diet of sampled individuals aligned well with previous isotope studies on diet preferences of southern stingrays (O’Shea et al., 2020; Tilley et al., 2013) but differed from diet estimates based on stomach content analyses. Stomach contents of southern stingrays from published studies show a higher proportion of teleosts and crustaceans, which comprised < 33 % of diet estimates in Bimini (Bigelow & Schroeder, 1953; Bowman et al., 2000; Gilliam & Sullivan, 1993; O’Shea et al., 2020; Randall, 1967; Smith & Herrnkind, 1992; Snelson Jr. & Williams, 1981). Stomach content analysis is, however, biased towards prey items with hard structures that are more difficult to digest (Baker et al., 2014; Hyslop, 1980), which could explain the greater importance of annelids in our models compared to assessments relying on stomach contents. Additionally, southern stingrays may only consume the soft body parts of molluscs (Randall, 1967), which could explain the higher proportion of molluscs in the diet when estimated using stable isotopes compared to stomach content analyses. We acknowledge, that estimating the diet via mixing models required the choice of stable isotope values of prey species described in studies from different study sites, and our isospace therefore may have limitations. Species-specific prey data from the study system are needed to refine future diet estimation.

### 4.2. Southern stingray condition

High trophic diversity as indicated by a range of δ^13^C values at the individual level could alternatively also result from habitat shifts following nutritional stress (Bowes et al., 2014; Catano et al., 2014; Hertz et al., 2015). However, an explanation for sex, spatial and seasonal differences in δ^13^C values based on nutritional state or condition is not supported by our results: across the four condition metrics we measured in serum samples of southern stingrays in Bimini, none indicated a proclivity for lowered condition or stress and we found no evidence for a correlation between δ^13^C values and serum condition markers.

Butyrate concentrations in this study were not different between seasons, locations, sex, or individuals of different sizes. This is surprising as varying butyrate concentrations between those variables have been previously described in other elasmobranchs and were suggested to be a consequence of physical activity, life stage and reproduction (Hammerschlag et al., 2018; Moorhead et al., 2021; Valls et al., 2016; Watson & Dickson, 2001). For example, in nurse sharks, *Ginglymostoma cirratum,* increased activity of males during mating season was hypothesized as a reason for lower butyrate values compared to females during the same period, and starvation during pregnancy was expected to increase butyrate concentrations in females during gestation periods (Moorhead et al., 2020; Wood et al., 2010). For southern stingrays, it is unclear if there are well defined mating and subsequent gestation periods around Bimini, and their reproductive cycle is thought to be asynchronous, i.e. parturition is expected to occur throughout the year (Ramirez Mosqueda et al., 2012). In agreement, we caught pregnant females year-round (Heim et al., unpublished data). Hence, when considering the gestation period of southern stingrays is < 6 months (Henningsen, 2000), and they can have ≥ 2 pregnancies a year (N. Mylniczenko, personal observation), gestation periods extended across our defined seasons potentially muting any reproductive effect. Consequently, we would not expect to find reproduction-related changes in butyrate concentrations between seasons. A total of six out of 122 sampled individuals showed elevated butyrate concentrations above 0.7 mmol/L. While these values still fell within the range of fed, fasted or post-captured elasmobranchs in wild and aquarium-managed scenarios (Speers-Roesch & Treberg, 2010; Dannemiller et al., 2023), they can also be indicative of 1) mobilised energy reserves, 2) low calories and 3) lipid depletion (Mylniczenko et al., unpublished data).

Only eight individuals in this study, seven of which were caught during summer, showed elevated concentrations of serum glucose, providing a very limited indication of stress. Generally, we found higher glucose concentrations in individuals sampled off South Bimini compared to Bonefish Hole and the lagoon during summer. This could be related to varying capture durations. Stingrays caught off South Bimini had easier access to deeper waters and escape routes. Once the rays were in deeper waters, the capture vessel had to spend more time guiding them towards the shallows and the mangroves once more, whereas individuals caught in the lagoon or in Bonefish Hole were more limited in their access to open water, so that captures were often shorter (V. Heim, personal observation). However, this could also be a physiological response to handling during the sampling process and related to observed patterns in lactate concentrations (see below).

While 26 individuals in this study showed serum lactate concentrations between 3 – 10 mmol/L indicative of muscle exertion, hypoventilation and stress, no individual experienced a physiological crisis (Mylniczenko et al., unpublished data). Previously, it was shown that lactate production in response to stress was reduced in larger elasmobranchs (Gallagher, Serafy, et al., 2014) but lactate concentrations were also shown to be positively correlated with the duration of the stress event and higher water temperatures in several species of sharks (Bouyoucos et al., 2023; Hoffmayer et al., 2012; Hyatt et al., 2018; Knotek et al., 2022). The increased serum lactate concentrations during the summer are thus likely a result of warmer water temperatures when the southern stingrays were caught and sampled. The increased lactate values in larger southern stingrays was likely a result of the duration of the sampling protocol, i.e. it was more difficult to restrain and control these individuals due to their increased strength and mass compared to smaller individuals and the subsequent increase in handling duration is expected to increase lactate concentrations (Bouyoucos et al., 2018; Fuller et al., 2020). This increase in lactate concentrations may also explain the observed decreasing trend in glucose concentrations with increasing disc widths tied to longer handling duration. Epaulette sharks, *Hemiscyllium ocellatum,* were shown to decrease glucose response and use ketone bodies to decrease lactate load during stress events (Dowd et al., 2010).

Beyond the physiological responses towards capture and handling, blood chemistry can also be informative for naturally induced stress. Hoffmayer et al. (2012) suggested that Atlantic sharpnose sharks, *Rhizoprionodon terranovae,* generally had higher blood glucose levels during seasons when the primary, i.e. neuroendocrine, stress response is increased. Among southern stingrays in Bimini, stress responses may be elevated during the winter due to an increased abundance of large apex predatory sharks such as great hammerheads and tiger sharks, and associated risk effects (Guttridge et al., 2017, 2022; Heim et al., 2021; Heithaus et al., 2008; Smukall et al., 2022). Furthermore, increased predator presence can lead to changes in the activity of prey (Cartamil et al., 2003, Bosiger & McCormick, 2014), which subsequently may impact serum butyrate concentrations (Watson & Dickson, 2001). However, despite expected seasonal and location specific predation risk butyrate concentrations were similar among animals sampled from Bonefish Hole, the lagoon and South Bimini during both seasons and we found no evidence for naturally induced stress in sampled southern stingrays. Furthermore, the hormone levels and blood gasses of southern stingrays measured during winter did not indicate an elevated stress response during this season (Wheaton et al. 2019). Additionally, animals in poor condition are known to engage more frequently in high-risk behaviours such as foraging in less safe habitats, so that the shallow water sampling might have biased our results towards animals in good condition that are more risk averse (Moran et al. 2021). We, however, acknowledge that most of our condition metrics reflect acute responses to stress given we lack methods to measure the effect of long-term, chronic stressors to fully understand potential effects of changing predator abundance on the condition of the southern stingrays.

Similar to the above physiological markers, we did not document osmolality measurements in serum that were indicative of stress, loss of liver condition or low urea levels. While we found a trend towards lower osmolality in females in the lagoon during winter compared to summer, fluctuations and seasonal changes of serum osmolality within the physiological range are common in marine teleosts and sharks (Hoffmayer et al., 2012; Mandrup-Poulsen, 1981; Sulikowski et al., 2003), and seasonal variation of osmolality has been suggested as a strategy in a variety of fish to reduce the cost of osmoregulation (Jobling, 1994). Thus, the observed variation in serum osmolality in female southern stingrays in the lagoon could be an indication that the females are using this strategy to reduce osmoregulatory costs, or may simply be a consequence of normal, physiological variation in osmolality given slightly changing osmolality (e.g. via precipitation, tidal stage, and evaporation; Newman et al., 2007) of the areas southern stingrays spend time in between the summer and winter. Consequently, the higher serum osmolality in the lagoon during summer may be a normal physiological response to increased evaporation due to higher temperatures in these very shallow water habitats. However, without accompanying water samples from when and where southern stingrays were caught, we cannot conclusively answer this question.

## 5. Conclusion

We used carbon and nitrogen stable isotopes to explore the trophic ecology of southern stingrays and provide measurements of four different serum metrics related to condition and stress responses in wild caught elasmobranchs. Our stable isotope data suggests that southern stingrays in Bimini are important mesopredators in seagrass and mangrove habitats with a preference for annelids and molluscs across all size classes and different seasons. Our measured serum concentrations of butyrates, glucose, lactate, and osmolality suggest that the southern stingrays sampled in this study were in overall good condition. However, the lack of changes in condition related serum concentrations in response to the tested variables, which were previously shown to alter the condition in other elasmobranch species, indicate that more work is needed to understand the suitability of assessed metrics as a non-lethal condition estimation in wild-caught southern stingrays. We thus suggest that future studies correlate measured serum concentrations of butyrates, glucose, lactate, and osmolality with ultrasonography data of the liver density (Greene et al., 2022), a reliable method to assess condition in elasmobranchs (Grant et al., 2013; Mylniczenko et al., 2018; Mylniczenko, 2012; Mylniczenko et al., 2017), to improve our understanding how blood chemical status relates to the overall health of these animals.

## Acknowledgement

Our gratitude goes to the Department of Fisheries and the Department of Marine Resources of The Bahamas for providing the necessary permits to conduct fieldwork in Bimini. All animals in this study were caught and all sampling protocols accepted under research permits MAMR/LIA/22 and and MA&MR/FIS/17B. Further, we wish to thank Cristina Mercoli and Daniel B. Nelson from the University of Basel for all their help during the stable isotope sample preparation and analysis of the muscle and serum samples at the Stable Isotope Ecology Lab in Basel. Thank you to Laura Garcia Barcia and Kirk Gastrich from the Florida International University for freeze-drying the muscle and serum samples prior to the stable isotope analysis. We are grateful to Catharine J. Wheaton for her help in analysing the serum samples for the condition metrics. We would like to thank Dieter Ebert and Valentin Amrhein from the University of Basel for their supervision of the first author during his PhD dissertation and for providing helpful comments during the development of this manuscript. We wish to thank Fränzi Korner-Nievergelt for her valuable inputs on the statistical methods. We are thankful to all the staff members and volunteers of the Bimini Biological Field Station Foundation for their help during southern stingray captures, sample processing and cataloguing. This study was supported by a grant of the Basler Stiftung für Experimentelle Zoologie, the Save Our Seas Foundation (grant #260), and the Bimini Biological Field Station Foundation (BBFSF).

## Author Contributions

Vital Heim: conceptualisation, data curation, formal analysis, investigation, methodology, project administration, resources, software, visualisation, writing - original draft, writing – review & editing. Matthew J. Smukall: funding acquisition, investigation, resources, writing – review & editing. Natalie D. Mylniczenko: data curation, investigation, resources, writing – review & editing. Charlene M. Burns: data curation, investigation, resources, writing – review & editing. Nigel E. Hussey: investigation, resources, writing – review & editing. Ansgar Kahmen: resources, writing – review & editing. Philip Matich: conceptualisation, methodology, supervision, writing – review & editing.

## Statements and Declarations

### Competing Interests

The authors herewith declare that there are no conflicts of interest.

## Data Availability Statement

Upon publication: The raw data on which calculations were performed and conclusions drawn are publicly available under: [INSERT LINK TO DATA REPOSITORY]. The accompanying code for the statistical analyses will be available in a GitHub repository: https://github.com/vheim/Heim-et-al_2025_JOURNAL.

## Declaration of Funding

This study was supported by a grant of the Basler Stiftung für Experimentelle Zoologie, the Save Our Seas Foundation (grant #260), and the Bimini Biological Field Station Foundation (BBFSF).

## Notes

### Competing Interest Statement

The authors have declared no competing interest.

