## Supplementary Material for "Seasonal dynamics in the trophic ecology and condition of a marine, benthic mesopredator, the southern stingray, *Hypanus americanus*"

*Supplementary Material and Methods*

To compare muscle samples analysed at the different universities using different lipid extraction protocols, we applied the lipid extraction protocol from the University of Windsor to a subset of 10 muscle samples already analysed at the University in Basel and repeated the analysis with the lipid extracted samples. We compared the $\delta^{13}C$ and $\delta^{15}N$ values between lipid-and-urea-extracted and urea-extracted-only samples and calculated a linear regression to adjust the values from the University of Basel using the resulting equation (see Supplementary Material, Fig. S1).

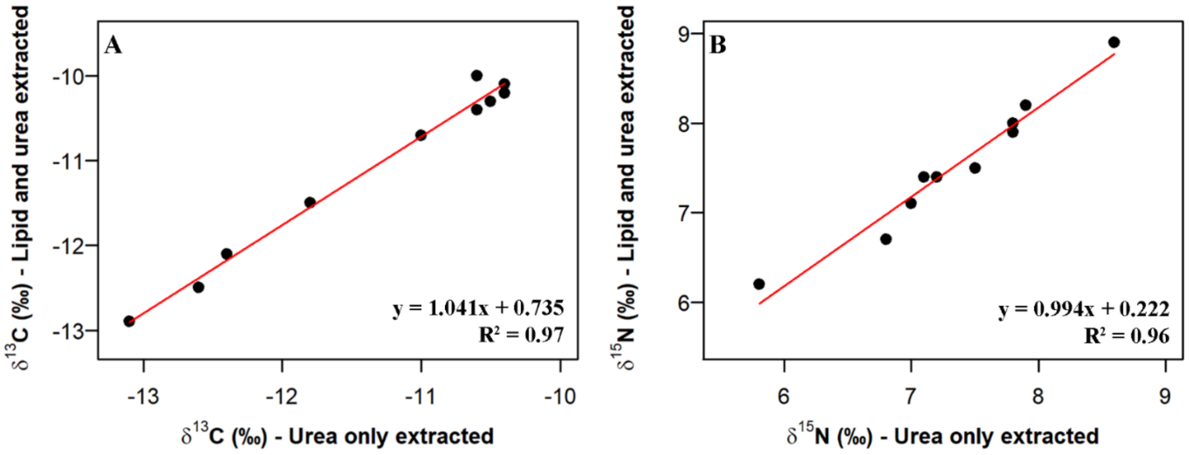


**Fig. S1. Comparison of (A) $\delta^{13}C$ and (B) $\delta^{15}N$ values in white muscle tissue samples where lipids and nitrogenous compounds have been extracted compared to white muscle tissue samples where only nitrogenous compounds were extracted.** The red line shows the linear regression line. The equation of the regression line and the $R^2$ value are shown in the bottom right corner of the figures. In the figures "urea" is used as a synonym for "nitrogenous compounds" for aesthetic reasons.

---

We used two-sample t-tests in a Bayesian framework in the *BayesFactor* R package (version 0.9.12-4.5; Morey & Rouder, 2023) to compare differences in the $\delta^{13}C$ and $\delta^{15}N$ values between samples analysed at different universities (i.e. Windsor and Basel), and between the two tissue types (i.e. muscle and serum samples). We sampled 2000 iterations and based statistical inference on the 95 % credible interval (95 % CrI) of the difference between the posterior means of the group, which was calculated using the *bayestestR* R package (version 0.17.0; Makowski et al., 2019).

*Supplementary Results*

For the analysis of stable isotope ratios in muscle samples, most samples were collected from mature females during summer (muscle samples: N = 70, serum samples: N = 64), but sample collection by season varied by sex and maturity state (see Supplementary Tables S1, S2), followed by mature females during winter (N = 32), immature females during summer (N = 24), mature males during winter (N = 11), immature females during winter and mature males during summer (N = 6 each).

**Table S1. Metadata of all southern stingrays sampled for white muscle tissue, including raw and diet-tissue discrimination factor (DTDF) corrected $\delta^{13}C$ and $\delta^{15}N$ values.** The PIT tag column contains the individual-specific number of the passive integrated transponder tag. PIT tag numbers in *italics* shows samples analysed at the University of Windsor, whereas all other PIT tags show samples analysed at the University of Basel. "F" in the Sex column shows females individuals, and "M" male individuals. The DW columns contains the disc width measurement in cm. The maturity column shows immature, mature and individuals of < 1 year of age (i.e. young-of-the-year ["yoy"]). The C:N column shows the ratio of carbon to nitrogen in each analysed sample. The columns $\delta^{13}C$ and $\delta^{15}N$ contain the raw isotopic ratios and the $\delta^{13}C$ DTDF and $\delta^{15}N$ DTDF contains the DTDF corrected values. The last column contains the capture location of the individuals (see Fig. 1) with the abbreviations: "LAG" – lagoon, "BFH" – Bonefish Hole, and "SB" – South Bimini.

| PIT tag | Date | Sex | DW *[cm]* | Maturity | C:N | $\delta^{13}C$ *[‰]* | $\delta^{13}C$ DTDF *[‰]* | $\delta^{15}N$ *[‰]* | $\delta^{15}N$ DTDF *[‰]* | Location |
|---|---|---|---|---|---|---|---|---|---|---|
| 989001026313501 | 2018-07-10 | F | 48.0 | immature | 3.3 | -11.1 | -12.8 | 7.1 | 3.4 | SB |
| 985121031803315 | 2018-07-11 | F | 77.0 | mature | 3.3 | -10.9 | -12.6 | 7.4 | 3.7 | SB |
| 989001004605331 | 2018-07-11 | F | 87.0 | mature | 3.2 | -10.3 | -12 | 8.8 | 5.1 | SB |
| 989001004378459 | 2018-07-12 | F | 99.0 | mature | 3.3 | -11.9 | -13.6 | 7.5 | 3.8 | SB |
| 989001005924984 | 2018-07-12 | F | 87.8 | mature | 3.3 | -11.8 | -13.5 | 7.7 | 4 | SB |
| 989001026313185 | 2018-07-12 | F | 52.0 | immature | 3.3 | -10.5 | -12.2 | 7.5 | 3.8 | SB |
| 989001026313143 | 2018-08-02 | F | 75.0 | mature | 3.2 | -12.2 | -13.9 | 7.5 | 3.8 | SB |

| | | | | | | | | | | |
|---|---|---|---|---|---|---|---|---|---|---|
| 989001026313517 | 2018-08-04 | F | 71.5 | immature | 3.3 | -12.4 | -14.1 | 8.3 | 4.6 | SB |
| 989001004605135 | 2018-08-02 | F | 90.0 | mature | 3.3 | -11.4 | -13.1 | 7.9 | 4.2 | SB |
| *989001026313424* | 2018-08-08 | F | 80.6 | mature | 3.2 | -10 | -11.7 | 6.8 | 3.1 | SB |
| *989001006700707* | 2018-08-09 | F | 67.6 | immature | 3 | -9.9 | -11.6 | 7.7 | 4 | SB |
| *989001005924838* | 2018-08-08 | F | 86.5 | mature | 3 | -9.9 | -11.6 | 6.9 | 3.2 | SB |
| *989001004605145* | 2018-10-01 | F | 91.6 | mature | 3.1 | -10.9 | -12.6 | 7.2 | 3.5 | SB |
| *989001006700794* | 2018-10-01 | F | 103.1 | mature | 3.1 | -10.9 | -12.6 | 7.1 | 3.4 | SB |
| *989001026313151* | 2018-10-01 | F | 77.0 | mature | 3.1 | -11.6 | -13.3 | 7.8 | 4.1 | SB |
| *989001006700633* | 2018-10-09 | F | 70.8 | immature | 3 | -11.6 | -13.3 | 6.2 | 2.5 | SB |
| *989001005924953* | 2018-10-28 | F | 103.2 | mature | 3.2 | -11.3 | -13 | 7.6 | 3.9 | SB |
| *989001005924862* | 2018-10-28 | F | 88.0 | mature | 3.3 | -11.7 | -13.4 | 7.7 | 4 | SB |
| *989001026313498* | 2018-10-28 | F | 85.2 | mature | 3.1 | -12.2 | -13.9 | 7.3 | 3.6 | SB |
| *989001006700659* | 2018-11-05 | F | 88.9 | mature | 3.1 | -9.9 | -11.6 | 7.7 | 4 | SB |
| *989001004378368* | 2018-11-08 | F | 101.2 | mature | 3.1 | -11.7 | -13.4 | 8.5 | 4.8 | SB |
| *989001004605300* | 2018-11-08 | F | 94.4 | mature | 3.2 | -12.2 | -13.9 | 8.3 | 4.6 | SB |
| *989001006700493* | 2018-11-08 | F | 78.1 | mature | 3.1 | -10.7 | -12.4 | 7.4 | 3.7 | SB |
| *985121031826185* | 2018-11-08 | F | 88.0 | mature | 3.2 | -13.7 | -15.4 | 7.6 | 3.9 | SB |
| *900236000107745* | 2018-11-15 | F | 98.8 | mature | 3.3 | -12.4 | -14.1 | 5.9 | 2.2 | SB |
| *989001006700845* | 2018-11-17 | F | 97.5 | mature | 3.2 | -12.1 | -13.8 | 7.6 | 3.9 | SB |
| *989001006700520* | 2018-11-17 | F | 94.0 | mature | 3.1 | -11.8 | -13.5 | 8.3 | 4.6 | SB |
| *989001026313302* | 2018-11-17 | F | 102.1 | mature | 3.2 | -11.7 | -13.4 | 7.5 | 3.8 | SB |
| *989001006700810* | 2018-11-17 | F | 88.3 | mature | 3.1 | -11.3 | -13 | 7.8 | 4.1 | SB |
| *989001026313537* | 2018-11-26 | F | 66.8 | immature | 3.1 | -9.8 | -11.5 | 8.9 | 5.2 | SB |
| *989001026313157* | 2018-11-26 | F | 84.0 | mature | 3.1 | -11.9 | -13.6 | 7.4 | 3.7 | SB |
| *989001026313413* | 2018-11-26 | F | 77.0 | mature | 3.2 | -12.5 | -14.2 | 7.5 | 3.8 | SB |
| *989001005924953* | 2018-12-01 | F | 98.9 | mature | 3.2 | -10.9 | -12.6 | 6.9 | 3.2 | SB |
| *989001026313275* | 2018-12-01 | F | 91.1 | mature | 3.2 | -11.2 | -12.9 | 7.5 | 3.8 | SB |
| *985121031809367* | 2018-12-01 | F | 96.0 | mature | 3.1 | -11.3 | -13 | 7.8 | 4.1 | SB |
| *989001026313529* | 2018-12-01 | F | 92.0 | mature | 3.1 | -10.6 | -12.3 | 8.3 | 4.6 | SB |
| *989001026313219* | 2018-12-01 | F | 87.2 | mature | 3.1 | -15.1 | -16.8 | 11.2 | 7.5 | SB |
| *989001026313492* | 2018-12-10 | F | 76.1 | mature | 3.1 | -10.6 | -12.3 | 7.4 | 3.7 | SB |
| *989001006700794* | 2019-01-25 | F | 95.0 | mature | 3 | -10.8 | -12.5 | 7.5 | 3.8 | SB |
| *989001026313278* | 2019-01-25 | F | 42.4 | immature | 3.1 | -11.6 | -13.3 | 8.7 | 5 | SB |
| *989001006700707* | 2019-01-25 | F | 67.1 | immature | 3 | -10.3 | -12 | 7.2 | 3.5 | SB |

| | | | | | | | | | | |
|---|---|---|---|---|---|---|---|---|---|---|
| *989001006700847* | 2019-01-25 | F | 91.1 | mature | 3.1 | -10.7 | -12.4 | 7.6 | 3.9 | SB |
| *985121031809367* | 2019-01-25 | F | 94.9 | mature | 3.1 | -11.2 | -12.9 | 8 | 4.3 | SB |
| *989001026313255* | 2019-02-06 | M | 50.0 | mature | 3.1 | -12.1 | -13.8 | 9.2 | 5.5 | SB |
| *989001004378515* | 2019-02-07 | F | 82.0 | mature | 3.2 | -11.4 | -13.1 | 7.9 | 4.2 | SB |
| *989001026313364* | 2019-02-15 | F | 86.8 | mature | 3.2 | -12.6 | -14.3 | 8.3 | 4.6 | SB |
| *989001004378356* | 2019-02-15 | F | 87.1 | mature | 3.2 | -10.1 | -11.8 | 8.1 | 4.4 | SB |
| *989001026313461* | 2019-02-15 | M | 51.2 | mature | 2.9 | -10.2 | -11.9 | 6.9 | 3.2 | SB |
| *989001026313160* | 2019-02-19 | M | 45.4 | mature | 3.2 | -11.8 | -13.5 | 8.5 | 4.8 | SB |
| *989001026313256* | 2019-02-19 | M | 46.0 | mature | 3.2 | -9.8 | -11.5 | 7.8 | 4.1 | SB |
| *989001026313322* | 2019-02-19 | M | 50.0 | mature | 3 | -9 | -10.7 | 8 | 4.3 | SB |
| *900236000107439* | 2019-02-24 | F | 91.0 | mature | 3.2 | -11.4 | -13.1 | 8.5 | 4.8 | SB |
| *989001026313189* | 2019-02-24 | M | 55.2 | mature | 3 | -8.8 | -10.5 | 8.2 | 4.5 | SB |
| *989001004378382* | 2019-02-04 | F | 82.4 | mature | 3.2 | -10.2 | -11.9 | 7.4 | 3.7 | SB |
| *989001006700661* | 2019-02-24 | F | 59.2 | immature | 3.2 | -12 | -13.7 | 8.1 | 4.4 | SB |
| *989001026313198* | 2019-02-24 | M | 42.9 | mature | 2.8 | -9.2 | -10.9 | 6.5 | 2.8 | SB |
| *900236000108035* | 2019-02-28 | F | 78.0 | mature | 3 | -11.9 | -13.6 | 6.4 | 2.7 | SB |
| *989001026313195* | 2019-02-28 | F | 42.0 | immature | 3 | -9.3 | -11 | 7.7 | 4 | SB |
| *989001005924973* | 2019-02-28 | F | 87.4 | mature | 3.1 | -12.5 | -14.2 | 7.8 | 4.1 | SB |
| *989001026313377* | 2019-02-28 | M | 48.0 | mature | 3.2 | -11 | -12.7 | 8.6 | 4.9 | SB |
| *985121031827394* | 2019-02-28 | F | 78.2 | mature | 3.2 | -13.3 | -15 | 6.8 | 3.1 | SB |
| *989001026313430* | 2019-03-03 | M | 49.6 | mature | 3.1 | -11.6 | -13.3 | 8.2 | 4.5 | SB |
| *989001026313172* | 2019-03-16 | F | 95.6 | mature | 3.1 | -13.4 | -15.1 | 7.2 | 3.5 | BFH |
| *989001026313388* | 2019-03-16 | F | 85.8 | mature | 3.1 | -10 | -11.7 | 7.1 | 3.4 | BFH |
| *989001026313378* | 2019-03-16 | F | 82.0 | mature | 3.2 | -12.2 | -13.9 | 7.2 | 3.5 | BFH |
| *989001026313357* | 2019-03-16 | F | 87.4 | mature | 3.2 | -11.2 | -12.9 | 6.3 | 2.6 | BFH |
| *989001005924856* | 2019-03-17 | F | 74.6 | immature | 3.1 | -12.9 | -14.6 | 6.8 | 3.1 | BFH |
| *989001026313343* | 2019-03-17 | F | 74.0 | immature | 3.2 | -12.6 | -14.3 | 7.6 | 3.9 | BFH |
| *989001026313359* | 2019-03-25 | F | 92.0 | mature | 3.1 | -13.7 | -15.4 | 7.4 | 3.7 | LAG |
| *989001026313370* | 2019-03-25 | M | 54.2 | mature | 3 | -10.8 | -12.5 | 7 | 3.3 | LAG |
| *989001026313187* | 2019-03-25 | F | 108.4 | mature | 3.4 | -13.1 | -14.8 | 8 | 4.3 | LAG |
| *989001026313440* | 2019-03-26 | M | 48.8 | mature | 3.1 | -11.4 | -13.1 | 7.5 | 3.8 | SB |
| *989001026313371* | 2019-03-26 | F | 83.0 | mature | 3.1 | -9.4 | -11.1 | 8.5 | 4.8 | SB |
| *985121031817057* | 2019-03-26 | F | 91.2 | mature | 3.2 | -11.2 | -12.9 | 7.8 | 4.1 | SB |
| *989001004378368* | 2019-04-20 | F | 94.0 | mature | 3.2 | -12.1 | -13.8 | 8.2 | 4.5 | SB |

| | | | | | | | | | | |
|---|---|---|---|---|---|---|---|---|---|---|
| *989001005924973* | 2019-04-20 | F | 92.0 | mature | 3.1 | -12.6 | -14.3 | 7.4 | 3.7 | SB |
| 989001026313417 | 2019-05-09 | F | 93.9 | mature | 3.3 | -13.2 | -14.9 | 7.3 | 3.6 | BFH |
| 989001026313369 | 2019-05-09 | F | 91.0 | mature | 3.3 | -12.1 | -13.8 | 6.1 | 2.4 | BFH |
| 989001026313152 | 2019-05-09 | F | 92.5 | mature | 3.1 | -12 | -13.7 | 7.5 | 3.8 | BFH |
| 989001026313194 | 2019-05-09 | F | 103.4 | mature | 3.2 | -12.4 | -14.1 | 7.6 | 3.9 | SB |
| 985121031803859 | 2019-05-09 | F | 88.3 | mature | 3.3 | -10.1 | -11.8 | 6.9 | 3.2 | BFH |
| 900236000108035 | 2019-06-28 | F | 74.5 | immature | 3.3 | -12.5 | -14.2 | 7.1 | 3.4 | SB |
| 989001026313493 | 2019-07-01 | F | 80.2 | mature | 3.3 | -11.9 | -13.6 | 8.1 | 4.4 | SB |
| 989001026313243 | 2019-07-01 | F | 86.0 | mature | 3.3 | -13.5 | -15.2 | 7.2 | 3.5 | SB |
| 989001026313369 | 2019-07-09 | F | 92.0 | mature | 3.2 | -12.4 | -14.1 | 6 | 2.3 | BFH |
| 989001006700560 | 2019-07-09 | F | 92.6 | mature | 3.3 | -10.5 | -12.2 | 7.6 | 3.9 | BFH |
| 989001026313193 | 2019-07-09 | F | 72.5 | immature | 3.3 | -9.5 | -11.2 | 7.5 | 3.8 | BFH |
| 989001026313538 | 2019-07-20 | F | 82.0 | mature | 3.3 | -14.1 | -15.8 | 6.6 | 2.9 | SB |
| 989001026313456 | 2019-07-20 | F | 98.0 | mature | 3.4 | -13.2 | -14.9 | 7.2 | 3.5 | LAG |
| 989001026313480 | 2019-07-20 | F | 69.5 | immature | 3.3 | -12.9 | -14.6 | 7.4 | 3.7 | LAG |
| 989001026313468 | 2019-07-22 | F | 83.0 | mature | 3.3 | -11.6 | -13.3 | 8 | 4.3 | LAG |
| 989001026313512 | 2019-07-22 | F | 96.0 | mature | 3.3 | -10.9 | -12.6 | 7.9 | 4.2 | LAG |
| 989001026313464 | 2019-07-22 | F | 105.6 | mature | 3.4 | -11.9 | -13.6 | 7.8 | 4.1 | SB |
| 989001026313449 | 2019-07-22 | F | 65.4 | immature | 3.2 | -11.1 | -12.8 | 7.5 | 3.8 | LAG |
| 900236000107745 | 2019-08-21 | F | 93.5 | mature | 3.1 | -10.7 | -12.4 | 7.2 | 3.5 | SB |
| 989001026313442 | 2019-08-21 | M | 42.6 | mature | 3.3 | -11.6 | -13.3 | 8.7 | 5 | SB |
| 989001026313467 | 2019-08-21 | F | 41.8 | yoy | 3.3 | -10.6 | -12.3 | 7.9 | 4.2 | SB |
| 989001026313503 | 2019-08-21 | M | 68.2 | mature | 3.3 | -11.3 | -13 | 9.7 | 6 | SB |
| 989001026313323 | 2019-08-21 | F | 41.9 | yoy | 3 | -7.9 | -9.6 | 6.7 | 3 | SB |
| 900236000107711 | 2019-09-12 | F | 103.0 | mature | 3.3 | -12.2 | -13.9 | 7.3 | 3.6 | SB |
| 989001006700537 | 2019-09-15 | F | 60.0 | immature | 3.3 | -10.5 | -12.2 | 8.1 | 4.4 | LAG |
| 989001026313532 | 2019-09-15 | F | 88.0 | mature | 3.2 | -10.2 | -11.9 | 8.5 | 4.8 | LAG |
| 989001026313424 | 2019-09-15 | F | 80.2 | mature | 3.3 | -9.9 | -11.6 | 8.3 | 4.6 | LAG |
| 989001026313473 | 2019-09-15 | F | 79.5 | mature | 4 | -13.2 | -14.9 | 8.1 | 4.4 | LAG |
| 989001026313463 | 2019-09-15 | F | 94.0 | mature | 3.2 | -10.2 | -11.9 | 8.6 | 4.9 | LAG |
| 989001030278495 | 2019-09-19 | F | 75.2 | mature | 3.3 | -12.1 | -13.8 | 7.9 | 4.2 | BFH |
| 989001030278548 | 2019-09-19 | F | 78.0 | mature | 3.2 | -11.8 | -13.5 | 7.3 | 3.6 | BFH |
| 989001030278436 | 2019-09-19 | F | 70.8 | immature | 3.3 | -9.6 | -11.3 | 7.4 | 3.7 | BFH |
| 989001030278568 | 2019-09-19 | F | 87.0 | mature | 3.3 | -12.5 | -14.2 | 7.3 | 3.6 | BFH |

| | | | | | | | | | | |
|---|---|---|---|---|---|---|---|---|---|---|
| 989001030278413 | 2019-09-19 | F | 83.9 | mature | 3.2 | -10.1 | -11.8 | 7 | 3.3 | BFH |
| 989001030278288 | 2019-09-19 | F | 89.2 | mature | 3.3 | -10.3 | -12 | 7.1 | 3.4 | BFH |
| 989001030278213 | 2019-09-19 | F | 85.5 | mature | 3.4 | -10.1 | -11.8 | 6.8 | 3.1 | BFH |
| 989001026313472 | 2019-09-20 | F | 89.0 | mature | 3.3 | -13.4 | -15.1 | 7.3 | 3.6 | BFH |
| 989001026313537 | 2019-09-24 | F | 69.5 | immature | 3.3 | -9.5 | -11.2 | 8.2 | 4.5 | SB |
| 989001026313488 | 2019-09-24 | F | 52.2 | immature | 3.3 | -10.5 | -12.2 | 8.7 | 5 | SB |
| 989001005924875 | 2019-09-24 | F | 92.1 | mature | 3.3 | -10.3 | -12 | 7.9 | 4.2 | SB |
| 989001004605295 | 2019-10-03 | F | 78.2 | mature | 3.3 | -11.8 | -13.5 | 7.6 | 3.9 | SB |
| 989001030278261 | 2019-10-04 | F | 83.0 | mature | 3.3 | -10.7 | -12.4 | 7.1 | 3.4 | BFH |
| 989001030278485 | 2019-10-04 | F | 70.4 | immature | 3.3 | -11.7 | -13.4 | 6.7 | 3 | BFH |
| 989001026313469 | 2019-10-04 | F | 101.0 | mature | 3.4 | -10.6 | -12.3 | 7.5 | 3.8 | BFH |
| 989001030278440 | 2019-10-04 | F | 91.5 | mature | 3.2 | -14.1 | -15.8 | 7.4 | 3.7 | BFH |
| 989001030278412 | 2019-10-04 | F | 90.4 | mature | 3.3 | -14.7 | -16.4 | 7.5 | 3.8 | BFH |
| 989001026313531 | 2019-10-04 | F | 95.5 | mature | 3.2 | -10.3 | -12 | 6.9 | 3.2 | BFH |
| 989001030278490 | 2019-10-05 | F | 78.0 | mature | 3.3 | -12.3 | -14 | 7.5 | 3.8 | LAG |
| 989001030278456 | 2019-10-05 | F | 80.3 | mature | 3.2 | -10.2 | -11.9 | 8 | 4.3 | LAG |
| 989001030278533 | 2019-10-05 | F | 73.0 | immature | 3.3 | -12.5 | -14.2 | 7.6 | 3.9 | LAG |
| 989001030278420 | 2019-10-05 | F | 68.3 | immature | 3.2 | -11.6 | -13.3 | 7.7 | 4 | LAG |
| 989001026313323 | 2019-10-13 | F | 39.6 | yoy | 3.3 | -9.6 | -11.3 | 7.6 | 3.9 | SB |
| 989001030278298 | 2019-10-13 | M | 45.0 | mature | 3.4 | -12.5 | -14.2 | 8.5 | 4.8 | SB |
| 989001030278227 | 2019-10-13 | F | 89.5 | mature | 3.3 | -12.1 | -13.8 | 8.6 | 4.9 | SB |
| 989001030278477 | 2019-10-21 | F | 45.0 | immature | 3.3 | -10.3 | -12 | 7.3 | 3.6 | SB |
| 989001030278461 | 2019-10-21 | M | 51.6 | mature | 3.3 | -11.8 | -13.5 | 8.4 | 4.7 | SB |
| 989001030278304 | 2019-11-01 | F | 72.5 | immature | 3.3 | -10.7 | -12.4 | 8.1 | 4.4 | LAG |
| 989001030278233 | 2019-11-01 | F | 91.7 | mature | 3.3 | -10.9 | -12.6 | 8.2 | 4.5 | LAG |
| 989001030278271 | 2019-11-01 | F | 78.4 | mature | 3.3 | -10.6 | -12.3 | 7.5 | 3.8 | LAG |
| 989001030278291 | 2019-11-01 | F | 88.5 | mature | 3.3 | -9.2 | -10.9 | 7.6 | 3.9 | LAG |
| 989001030278237 | 2019-11-01 | F | 78.5 | mature | 3.3 | -10.9 | -12.6 | 7.6 | 3.9 | LAG |
| 989001026313180 | 2019-11-01 | F | 77.0 | mature | 3.2 | -10.1 | -11.8 | 8 | 4.3 | LAG |
| 989001030278434 | 2019-11-01 | M | 51.9 | mature | 3.4 | -9.3 | -11 | 7.9 | 4.2 | LAG |
| 989001030278429 | 2019-11-01 | F | 90.5 | mature | 3.3 | -10 | -11.7 | 7.7 | 4 | LAG |
| 989001030278549 | 2019-11-01 | F | 73.2 | immature | 3.3 | -9.9 | -11.6 | 8.2 | 4.5 | LAG |
| 989001030278216 | 2019-11-01 | F | 85.7 | mature | 3.4 | -11.4 | -13.1 | 7.9 | 4.2 | LAG |
| 989001006700707 | 2019-11-12 | F | 71.2 | immature | 3.3 | -12.1 | -13.8 | 7.1 | 3.4 | SB |

| | | | | | | | | | | |
|---|---|---|---|---|---|---|---|---|---|---|
| 989001030278310 | 2019-11-12 | F | 52.0 | immature | 3.9 | -14.2 | -15.9 | 8.7 | 5 | SB |
| 989001030278219 | 2019-11-12 | F | 63.5 | immature | 3.2 | -11.3 | -13 | 8.6 | 4.9 | SB |
| 989001030278446 | 2019-11-12 | M | 53.0 | mature | 3.3 | -10.4 | -12.1 | 9.1 | 5.4 | SB |
| 989001030278567 | 2019-11-12 | F | 71.0 | immature | 3.2 | -9.3 | -11 | 7.2 | 3.5 | SB |
| 989001030278551 | 2019-11-25 | F | 86.7 | mature | 3.3 | -12.6 | -14.3 | 7.2 | 3.5 | BFH |
| 989001030278599 | 2019-11-25 | F | 92.7 | mature | 3.3 | -13.3 | -15 | 7.5 | 3.8 | BFH |
| 989001030278545 | 2019-11-25 | F | 99.5 | mature | 3.3 | -12.5 | -14.2 | 7.1 | 3.4 | BFH |
| 989001030278435 | 2019-11-27 | F | 89.0 | mature | 3.3 | -12.9 | -14.6 | 7.8 | 4.1 | BFH |
| 989001030278562 | 2019-11-27 | F | 97.9 | mature | 3.3 | -15 | -16.7 | 7.6 | 3.9 | BFH |

For the analysis of stable isotope ratios in serum samples, we collected 64 serum samples for mature female southern stingrays during summer and 38 during winter. A total of 24 serum samples of immature females were collected during summer and four during winter. We sampled 12 mature males for serum, of which eight samples were collected during summer and four samples during winter.

**Table S2. Metadata of all southern stingrays sampled for blood, including raw and diet-tissue discrimination factor (DTDF) corrected $\delta^{13}C$ and $\delta^{15}N$ values.** The PIT tag column contains the individual-specific number of the passive integrated transponder tag. "F" in the Sex column shows females individuals, and "M" male individuals. The DW columns contains the disc width measurement in cm. The maturity column shows immature, mature and individuals of < 1 year of age (i.e. young-of-the-year ["yoy"]). The C:N column shows the ratio of carbon to nitrogen in each analysed sample. The columns $\delta^{13}C$ and $\delta^{15}N$ contain the raw isotopic ratios and the $\delta^{13}C$ DTDF and $\delta^{15}N$ DTDF contains the DTDF corrected values. The last column contains the capture location of the individuals (see Fig. 1) with the abbreviations: "LAG" – lagoon, "BFH" – Bonefish Hole, and "SB" – South Bimini.

| PIT tag | Date | Sex | DW *[cm]* | Maturity | C:N | $\delta^{13}C$ *[‰]* | $\delta^{13}C$ DTDF *[‰]* | $\delta^{15}N$ *[‰]* | $\delta^{15}N$ DTDF *[‰]* | Location |
|---|---|---|---|---|---|---|---|---|---|---|
| 989001026313501 | 2018-07-10 | F | 48.0 | immature | 4.1 | -12.6 | -15.4 | 5.2 | 3.0 | SB |
| 989001026313238 | 2018-07-10 | F | 77.2 | mature | 3.5 | -11.5 | -14.3 | 7.1 | 4.9 | SB |
| 985121031803315 | 2018-07-11 | F | 77.0 | mature | 3.8 | -12.0 | -14.8 | 6.1 | 3.9 | SB |
| 989001004605331 | 2018-07-11 | F | 87.0 | mature | 3.6 | -11.7 | -14.5 | 6.3 | 4.1 | SB |
| 989001026313185 | 2018-07-12 | F | 52.0 | immature | 4.1 | -13.2 | -16.0 | 5.7 | 3.5 | SB |
| 989001006700866 | 2018-07-12 | M | 57.0 | mature | 3.7 | -10.8 | -13.6 | 8.0 | 5.8 | SB |
| 989001004605180 | 2018-07-12 | F | 68.0 | immature | 3.9 | -14.7 | -17.5 | 4.2 | 2.0 | SB |
| 989001005924984 | 2018-07-12 | F | 87.8 | mature | 3.6 | -13.3 | -16.1 | 6.2 | 4.0 | SB |
| 989001004378459 | 2018-07-12 | F | 99.0 | mature | 3.8 | -13.5 | -16.3 | 5.9 | 3.7 | SB |
| 989001005924837 | 2018-08-02 | F | 98.2 | mature | 3.7 | -11.4 | -14.2 | 5.5 | 3.3 | SB |
| 989001004605135 | 2018-08-02 | F | 90.0 | mature | 3.9 | -12.9 | -15.7 | 6.0 | 3.8 | SB |
| 989001026313458 | 2018-08-05 | F | 93.3 | mature | 3.7 | -12.3 | -15.1 | 5.2 | 3.0 | SB |
| 989001004605145 | 2018-10-01 | F | 91.6 | mature | 3.9 | -12.1 | -14.9 | 5.7 | 3.5 | SB |
| 989001006700794 | 2018-10-01 | F | 103.1 | mature | 3.7 | -12.4 | -15.2 | 5.5 | 3.3 | SB |
| 989001006700633 | 2018-10-09 | F | 70.6 | immature | 3.5 | -15.8 | -18.6 | 3.6 | 1.4 | SB |

| | | | | | | | | | | |
|---|---|---|---|---|---|---|---|---|---|---|
| 989001026313498 | 2018-10-28 | F | 85.2 | mature | 3.3 | -12.6 | -15.4 | 5.3 | 3.1 | SB |
| 989001026313524 | 2018-11-03 | F | 93.1 | mature | 3.6 | -13.0 | -15.8 | 5.3 | 3.1 | SB |
| 989001006700659 | 2018-11-05 | F | 88.9 | mature | 3.7 | -10.9 | -13.7 | 5.4 | 3.2 | SB |
| 989001006700493 | 2018-11-08 | F | 78.1 | mature | 3.5 | -11.5 | -14.3 | 5.1 | 2.9 | SB |
| 989001004605300 | 2018-11-08 | F | 94.4 | mature | 3.7 | -12.3 | -15.1 | 6.0 | 3.8 | SB |
| 989001004378368 | 2018-11-08 | F | 101.2 | mature | 3.5 | -12.5 | -15.3 | 5.7 | 3.5 | SB |
| 989001005924813 | 2018-11-17 | F | 65.2 | immature | 3.5 | -11.5 | -14.3 | 5.1 | 2.9 | SB |
| 989001026313302 | 2018-11-17 | F | 102.1 | mature | 4.0 | -12.2 | -15.0 | 6.3 | 4.1 | SB |
| 989001004605145 | 2018-11-17 | F | 90.1 | mature | 3.8 | -12.3 | -15.1 | 5.7 | 3.5 | SB |
| 989001006700520 | 2018-11-17 | F | 94.0 | mature | 3.9 | -13.4 | -16.2 | 5.5 | 3.3 | SB |
| 989001005924837 | 2018-11-17 | F | 104.8 | mature | 3.6 | -11.4 | -14.2 | 5.6 | 3.4 | SB |
| 989001026313143 | 2018-11-17 | F | 77.0 | mature | 3.8 | -15.0 | -17.8 | 5.1 | 2.9 | SB |
| 989001006700810 | 2018-11-17 | F | 88.3 | mature | 4.1 | -13.0 | -15.8 | 5.5 | 3.3 | SB |
| 989001026313157 | 2018-11-26 | F | 84.0 | mature | 3.8 | -14.1 | -16.9 | 5.1 | 2.9 | SB |
| 989001026313413 | 2018-11-26 | F | 77.0 | mature | 3.6 | -12.5 | -15.3 | 5.2 | 3.0 | SB |
| 989001026313537 | 2018-11-26 | F | 66.8 | immature | 3.8 | -9.7 | -12.5 | 7.2 | 5.0 | SB |
| 989001005924851 | 2018-11-26 | F | 79.4 | mature | 3.8 | -12.7 | -15.5 | 4.7 | 2.5 | SB |
| 989001026313529 | 2018-12-01 | F | 92.0 | mature | 3.8 | -11.8 | -14.6 | 6.0 | 3.8 | SB |
| 989001004605331 | 2018-12-01 | F | 99.4 | mature | 3.9 | -12.3 | -15.1 | 6.4 | 4.2 | SB |
| 989001026313219 | 2018-12-01 | F | 87.2 | mature | 4.1 | -16.0 | -18.8 | 6.5 | 4.3 | SB |
| 989001026313275 | 2018-12-01 | F | 91.1 | mature | 3.8 | -12.6 | -15.4 | 5.0 | 2.8 | SB |
| 900236000107745 | 2018-12-01 | F | 98.8 | mature | 4.0 | -13.7 | -16.5 | 5.7 | 3.5 | SB |
| 985121031809367 | 2018-12-01 | F | 96.0 | mature | 3.7 | -12.7 | -15.5 | 6.2 | 4.0 | SB |
| 989001006700810 | 2018-12-10 | F | 88.3 | mature | 4.0 | -13.0 | -15.8 | 5.6 | 3.4 | SB |
| 989001004605145 | 2018-12-10 | F | 93.0 | mature | 3.9 | -12.4 | -15.2 | 5.7 | 3.5 | SB |
| 989001026313492 | 2018-12-10 | F | 76.1 | mature | 3.5 | -11.6 | -14.4 | 5.4 | 3.2 | SB |
| NA | 2018-12-10 | F | 96.4 | mature | 3.8 | -11.6 | -14.4 | 6.8 | 4.6 | SB |
| 985121031809367 | 2019-01-25 | F | 94.9 | mature | 3.7 | -12.4 | -15.2 | 6.6 | 4.4 | SB |
| 989001006700707 | 2019-01-25 | F | 67.1 | immature | 3.6 | -14.6 | -17.4 | 4.3 | 2.1 | SB |
| 989001026313278 | 2019-01-25 | M | 42.2 | mature | 3.7 | -12.2 | -15.0 | 7.7 | 5.5 | SB |
| 989001006700794 | 2019-01-25 | F | 95.0 | mature | 3.7 | -12.3 | -15.1 | 6.0 | 3.8 | SB |
| 989001006700847 | 2019-01-25 | F | 91.1 | mature | 3.8 | -12.0 | -14.8 | 6.1 | 3.9 | SB |
| 989001004378382 | 2019-02-04 | F | 82.4 | mature | 3.8 | -11.4 | -14.2 | 5.7 | 3.5 | SB |
| 989001026313255 | 2019-02-06 | M | 50.0 | mature | 3.6 | -13.1 | -15.9 | 8.4 | 6.2 | SB |

| | | | | | | | | | | |
|---|---|---|---|---|---|---|---|---|---|---|
| 989001006700847 | 2019-02-07 | F | 94.9 | mature | 3.8 | -12.6 | -15.4 | 5.8 | 3.6 | SB |
| 989001004378515 | 2019-02-07 | F | 82.0 | mature | 3.9 | -12.9 | -15.7 | 5.9 | 3.7 | SB |
| 989001026313200 | 2019-02-07 | F | 91.8 | mature | 3.7 | -10.0 | -12.8 | 5.3 | 3.1 | SB |
| 989001026313357 | 2019-03-16 | F | 87.4 | mature | 3.5 | -12.6 | -15.4 | 4.5 | 2.3 | BFH |
| 989001026313388 | 2019-03-16 | F | 85.8 | mature | 3.6 | -11.1 | -13.9 | 4.6 | 2.4 | BFH |
| 989001026313378 | 2019-03-16 | F | 82.0 | mature | 3.4 | -14.6 | -17.4 | 4.3 | 2.1 | BFH |
| 989001026313172 | 2019-03-16 | F | 95.6 | mature | 4.0 | -15.8 | -18.6 | 4.7 | 2.5 | BFH |
| 989001005924856 | 2019-03-17 | F | 74.6 | immature | 3.8 | -13.6 | -16.4 | 4.1 | 1.9 | BFH |
| 989001026313343 | 2019-03-17 | F | 74.0 | immature | 3.8 | -13.7 | -16.5 | 4.2 | 2.0 | BFH |
| 989001026313402 | 2019-03-25 | F | 88.5 | mature | 3.5 | -14.2 | -17.0 | 6.2 | 4.0 | LAG |
| 989001026313187 | 2019-03-25 | F | 108.4 | mature | 3.6 | -14.7 | -17.5 | 5.8 | 3.6 | LAG |
| 989001026313370 | 2019-03-25 | M | 54.2 | mature | 3.8 | -12.6 | -15.4 | 5.8 | 3.6 | LAG |
| 989001026313359 | 2019-03-25 | F | 92.0 | mature | 3.6 | -15.6 | -18.4 | 5.7 | 3.5 | LAG |
| NA | 2019-03-26 | F | 84.0 | mature | 3.6 | -16.8 | -19.6 | 9.4 | 7.2 | SB |
| 989001005924975 | 2019-03-26 | F | 97.4 | mature | 3.5 | -13.0 | -15.8 | 6.2 | 4.0 | SB |
| 985121031817057 | 2019-03-26 | F | 91.2 | mature | 3.7 | -12.4 | -15.2 | 6.2 | 4.0 | SB |
| 989001006700794 | 2019-03-26 | F | 97.0 | mature | 3.6 | -12.0 | -14.8 | 6.2 | 4.0 | SB |
| 989001026313197 | 2019-03-26 | F | 72.6 | immature | 3.3 | -12.9 | -15.7 | 7.1 | 4.9 | SB |
| 989001026313440 | 2019-03-26 | M | 48.8 | mature | 3.5 | -13.6 | -16.4 | 6.9 | 4.7 | SB |
| 989001026313371 | 2019-03-26 | F | 83.0 | mature | 3.5 | -9.9 | -12.7 | 7.1 | 4.9 | SB |
| 989001004378368 | 2019-04-20 | F | 94.0 | mature | 3.8 | -13.1 | -15.9 | 6.4 | 4.2 | SB |
| 989001026313417 | 2019-05-09 | F | 93.9 | mature | 3.8 | -14.6 | -17.4 | 5.0 | 2.8 | BFH |
| 989001026313369 | 2019-05-09 | F | 91.0 | mature | 3.8 | -13.5 | -16.3 | 4.4 | 2.2 | BFH |
| 989001026313152 | 2019-05-09 | F | 92.5 | mature | 3.5 | -14.6 | -17.4 | 4.2 | 2.0 | BFH |
| 989001026313194 | 2019-05-09 | F | 103.4 | mature | 3.8 | -13.8 | -16.6 | 5.2 | 3.0 | SB |
| 989001026313287 | 2019-05-09 | F | 93.1 | mature | 4.0 | -14.2 | -17.0 | 4.7 | 2.5 | BFH |
| 989001026313486 | 2019-05-09 | F | 88.0 | mature | 3.6 | -12.6 | -15.4 | 4.7 | 2.5 | BFH |
| 989001026313525 | 2019-05-09 | F | 90.3 | mature | 3.7 | -13.2 | -16.0 | 4.1 | 1.9 | BFH |
| 985121031803859 | 2019-05-09 | F | 88.3 | mature | 3.8 | -12.2 | -15.0 | 4.5 | 2.3 | BFH |
| 900236000108035 | 2019-06-28 | F | 74.5 | immature | 4.0 | -14.0 | -16.8 | 5.0 | 2.8 | SB |
| 989001026313243 | 2019-07-01 | F | 86.0 | mature | 4.1 | -15.2 | -18.0 | 4.8 | 2.6 | SB |
| 989001026313198 | 2019-07-01 | M | 45.0 | mature | 3.9 | -11.9 | -14.7 | 6.0 | 3.8 | SB |
| 989001026313493 | 2019-07-01 | F | 80.2 | mature | 3.5 | -15.0 | -17.8 | 5.2 | 3.0 | SB |
| 989001006700567 | 2019-07-01 | F | 86.5 | mature | 3.8 | -12.3 | -15.1 | 5.4 | 3.2 | SB |

| | | | | | | | | | | |
|---|---|---|---|---|---|---|---|---|---|---|
| 989001006700560 | 2019-07-09 | F | 92.6 | mature | 3.7 | -12.0 | -14.8 | 5.1 | 2.9 | BFH |
| 989001026313369 | 2019-07-09 | F | 92.0 | mature | 4.2 | -14.4 | -17.2 | 3.8 | 1.6 | BFH |
| 989001026313510 | 2019-07-09 | F | 89.0 | mature | 4.0 | -14.1 | -16.9 | 4.4 | 2.2 | BFH |
| 989001026313193 | 2019-07-09 | F | 72.5 | immature | 3.5 | -11.4 | -14.2 | 5.7 | 3.5 | BFH |
| 989001026313538 | 2019-07-20 | F | 82.0 | mature | 4.3 | -16.6 | -19.4 | 3.6 | 1.4 | SB |
| 989001026313456 | 2019-07-20 | F | 98.0 | mature | 3.9 | -14.7 | -17.5 | 4.7 | 2.5 | LAG |
| 989001026313449 | 2019-07-22 | F | 65.4 | immature | 4.2 | -12.8 | -15.6 | 5.2 | 3.0 | LAG |
| 989001026313468 | 2019-07-22 | F | 83.0 | mature | 3.8 | -12.3 | -15.1 | 5.4 | 3.2 | LAG |
| 989001026313464 | 2019-07-22 | F | 105.6 | mature | 4.4 | -13.6 | -16.4 | 6.1 | 3.9 | SB |
| 989001026313161 | 2019-08-13 | F | 56.4 | immature | 4.0 | -12.4 | -15.2 | 5.9 | 3.7 | SB |
| 989001004378459 | 2019-08-21 | F | 93.7 | mature | 4.1 | -14.3 | -17.1 | 5.6 | 3.4 | SB |
| 989001026313444 | 2019-08-21 | F | 53.5 | immature | 4.3 | -11.9 | -14.7 | 7.2 | 5.0 | SB |
| 989001026313467 | 2019-08-21 | F | 41.8 | yoy | 3.7 | -12.3 | -15.1 | 7.0 | 4.8 | SB |
| 989001026313514 | 2019-08-21 | F | 46.3 | immature | 4.2 | -12.3 | -15.1 | 6.5 | 4.3 | SB |
| 900236000107745 | 2019-08-21 | F | 93.5 | mature | 4.2 | -15.0 | -17.8 | 5.0 | 2.8 | SB |
| 989001026313503 | 2019-08-21 | M | 48.2 | mature | 3.8 | -12.1 | -14.9 | 7.8 | 5.6 | SB |
| 989001026313442 | 2019-08-21 | M | 42.6 | mature | 4.0 | -12.2 | -15.0 | 7.3 | 5.1 | SB |
| 989001026313473 | 2019-09-15 | F | 79.5 | mature | 4.0 | -11.2 | -14.0 | 6.1 | 3.9 | LAG |
| 989001006700537 | 2019-09-15 | F | 60.0 | immature | 3.9 | -12.3 | -15.1 | 5.7 | 3.5 | LAG |
| 989001026313424 | 2019-09-15 | F | 80.2 | mature | 3.9 | -11.1 | -13.9 | 6.0 | 3.8 | LAG |
| 989001026313495 | 2019-09-15 | F | 76.5 | mature | 3.9 | -11.4 | -14.2 | 5.4 | 3.2 | LAG |
| 989001026313532 | 2019-09-15 | F | 88.0 | mature | 3.7 | -11.3 | -14.1 | 5.9 | 3.7 | LAG |
| 989001030278495 | 2019-09-19 | F | 75.2 | mature | 4.0 | -12.2 | -15.0 | 5.6 | 3.4 | BFH |
| 989001030278436 | 2019-09-19 | F | 70.8 | immature | 3.9 | -11.1 | -13.9 | 4.8 | 2.6 | BFH |
| 989001030278548 | 2019-09-19 | F | 78.0 | mature | 4.0 | -13.6 | -16.4 | 4.4 | 2.2 | BFH |
| 989001030278413 | 2019-09-19 | F | 83.9 | mature | 3.7 | -11.9 | -14.7 | 4.8 | 2.6 | BFH |
| 989001026313488 | 2019-09-24 | F | 54.2 | immature | 4.2 | -12.6 | -15.4 | 6.3 | 4.1 | SB |
| 989001005924875 | 2019-09-24 | F | 92.1 | mature | 3.7 | -11.9 | -14.7 | 6.0 | 3.8 | SB |
| 989001030278412 | 2019-10-04 | F | 90.4 | mature | 3.7 | -15.8 | -18.6 | 5.2 | 3.0 | BFH |
| 989001026313469 | 2019-10-04 | F | 101.0 | mature | 3.9 | -12.2 | -15.0 | 4.8 | 2.6 | BFH |
| 989001030278485 | 2019-10-04 | F | 70.4 | immature | 4.0 | -14.6 | -17.4 | 4.1 | 1.9 | BFH |
| 989001030278261 | 2019-10-04 | F | 83.0 | mature | 3.7 | -12.2 | -15.0 | 4.8 | 2.6 | BFH |
| 989001030278440 | 2019-10-04 | F | 91.5 | mature | 4.3 | -15.8 | -18.6 | 4.9 | 2.7 | BFH |
| 989001030278456 | 2019-10-05 | F | 80.3 | mature | 3.9 | -11.3 | -14.1 | 5.9 | 3.7 | LAG |

| | | | | | | | | | | |
|---|---|---|---|---|---|---|---|---|---|---|
| 989001030278533 | 2019-10-05 | F | 73.0 | immature | 4.1 | -13.5 | -16.3 | 5.7 | 3.5 | LAG |
| 989001030278490 | 2019-10-05 | F | 78.0 | mature | 3.9 | -13.2 | -16.0 | 5.7 | 3.5 | LAG |
| 989001030278420 | 2019-10-05 | F | 68.3 | immature | 3.8 | -12.0 | -14.8 | 5.1 | 2.9 | LAG |
| 989001026313323 | 2019-10-13 | F | 39.6 | yoy | 3.7 | -11.1 | -13.9 | 5.8 | 3.6 | SB |
| 989001030278298 | 2019-10-13 | M | 45.0 | mature | 3.8 | -13.9 | -16.7 | 7.8 | 5.6 | SB |
| 989001026313461 | 2019-10-21 | M | 51.6 | mature | 3.8 | -12.1 | -14.9 | 7.1 | 4.9 | SB |
| 989001030278610 | 2019-10-21 | F | 64.0 | immature | 3.7 | -12.0 | -14.8 | 6.6 | 4.4 | SB |
| 989001030278429 | 2019-11-01 | F | 90.5 | mature | 3.7 | -11.2 | -14.0 | 5.8 | 3.6 | LAG |
| 989001030278304 | 2019-11-01 | F | 72.5 | immature | 3.9 | -11.8 | -14.6 | 6.0 | 3.8 | LAG |
| 989001030278216 | 2019-11-01 | F | 85.7 | mature | 3.8 | -12.3 | -15.1 | 5.8 | 3.6 | LAG |
| 989001030278233 | 2019-11-01 | F | 91.7 | mature | 3.7 | -12.5 | -15.3 | 6.3 | 4.1 | LAG |
| 989001026313180 | 2019-11-01 | F | 77.0 | mature | 3.8 | -11.7 | -14.5 | 6.1 | 3.9 | LAG |
| 989001030278434 | 2019-11-01 | M | 51.9 | mature | 3.5 | -9.9 | -12.7 | 6.0 | 3.8 | LAG |
| 989001030278549 | 2019-11-01 | F | 73.2 | immature | 3.7 | -10.4 | -13.2 | 5.8 | 3.6 | LAG |
| 989001030278291 | 2019-11-01 | F | 88.5 | mature | 3.5 | -10.0 | -12.8 | 5.5 | 3.3 | LAG |
| 989001030278271 | 2019-11-01 | F | 78.4 | mature | 3.7 | -11.5 | -14.3 | 5.2 | 3.0 | LAG |
| 989001030278237 | 2019-11-01 | F | 78.5 | mature | 4.1 | -12.2 | -15.0 | 5.1 | 2.9 | LAG |
| 989001030278567 | 2019-11-12 | F | 71.0 | immature | 3.7 | -10.8 | -13.6 | 6.3 | 4.1 | SB |
| 989001030278219 | 2019-11-12 | F | 63.5 | immature | 3.7 | -13.0 | -15.8 | 6.6 | 4.4 | SB |
| 989001030278446 | 2019-11-12 | M | 53.0 | mature | 3.4 | -11.0 | -13.8 | 8.7 | 6.5 | SB |
| 989001030278310 | 2019-11-12 | F | 52.0 | immature | 4.3 | -11.3 | -14.1 | 7.0 | 4.8 | SB |
| 989001030278545 | 2019-11-25 | F | 99.5 | mature | 3.6 | -14.4 | -17.2 | 5.2 | 3.0 | BFH |
| 989001030278551 | 2019-11-25 | F | 86.1 | mature | 3.9 | -14.9 | -17.7 | 4.7 | 2.5 | BFH |
| 989001030278599 | 2019-11-25 | F | 92.7 | mature | 3.8 | -14.4 | -17.2 | 5.6 | 3.4 | BFH |
| 989001030278435 | 2019-11-27 | F | 89.0 | mature | 3.8 | -14.4 | -17.2 | 5.4 | 3.2 | BFH |
| 989001030278562 | 2019-11-27 | F | 97.9 | mature | 4.1 | -17.0 | -19.8 | 5.4 | 3.2 | BFH |
| 989001030278265 | 2019-11-27 | F | 84.8 | mature | 3.9 | -12.7 | -15.5 | 4.4 | 2.2 | BFH |

Mean isotopic ratios from muscle samples analysed at the University of Windsor ($\delta^{13}C$ : -11.4 ± 1.2 ‰, $\delta^{15}N$:  4.0 ± 0.8, N = 67) and at the University of Basel ($\delta^{13}C$ : -13.2 ± 1.3 ‰, $\delta^{15}N$: 4.0 ± 0.6, N = 82) did not differ ($\delta^{13}C$: 95 % CrI [-0.40 ‰, 0.21 ‰], $\delta^{15}N$: 95 % CrI [-0.30 ‰, 0.31 ‰], Fig. S2). Serum $\delta^{13}C$ and $\delta^{15}N$ values were lower than corresponding values in muscle tissue ($\delta^{13}C$: 95 % CrI [-2.07 ‰, -1.52 ‰], $\delta^{15}N$: 95 % CrI [-0.82 ‰, -0.35 ‰]).

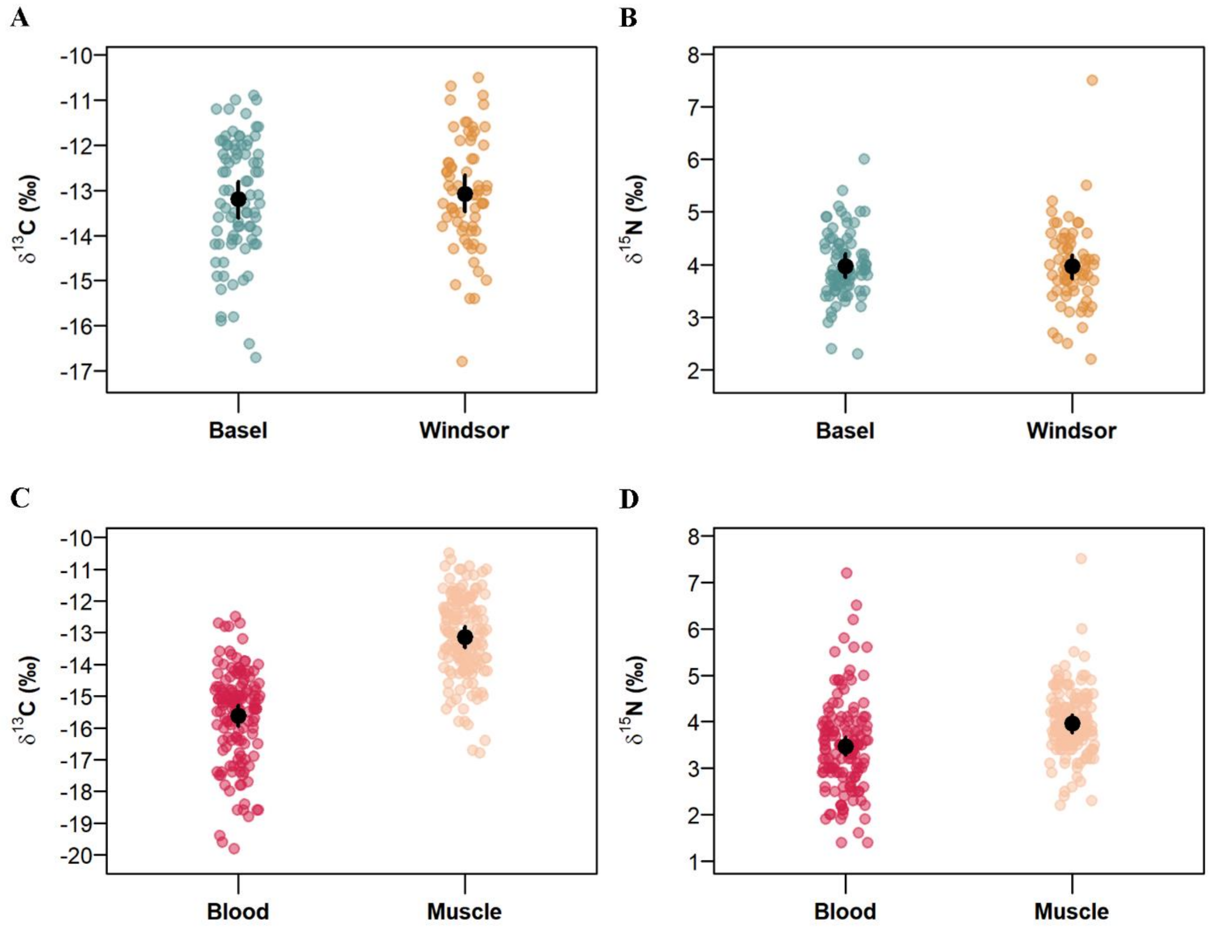


**Fig. S2. Comparison of (A) $\delta^{13}C$ and (B) $\delta^{15}N$ values of white muscle tissue samples analysed at the University of Windsor and the University of Basel, and comparison of (C) $\delta^{13}C$ and (D) $\delta^{15}N$ values from white muscle tissue and serum, i.e. blood, samples.** The coloured dots show diet-tissue discrimination factor (DTDF) corrected values. The black dots show the mean isotopic ratios per group.

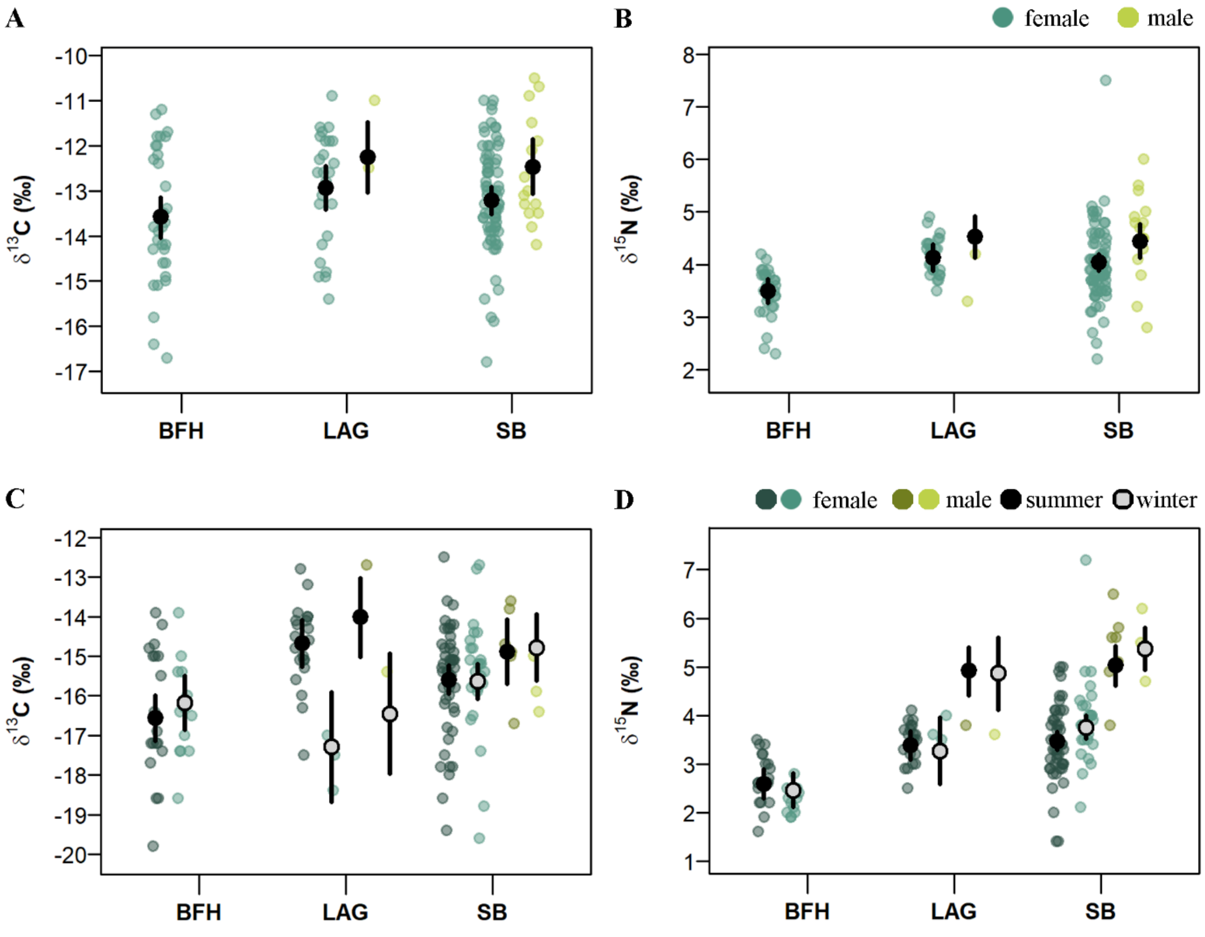


**Fig. S3. Effects of sex and location on the (A) $\delta^{13}C$ and (B) $\delta^{15}N$ values of white muscle tissue samples, and effects of sex, season and location on the (C) $\delta^{13}C$ and (B) $\delta^{15}N$ values of serum samples.** Coloured points show the raw isotopic ratios by sex and season. Isotopic values from serum samples in panels C and D are shown for the two seasons summer (darker colours) and winter (lighter colours). Capture locations are abbreviated as "LAG" – lagoon, "BFH" – Bonefish Hole, and "SB" – South Bimini. Posterior means simulated from the liner mixed models (LMMs) are shown by black (summer) and grey points (winter) and 95 % credible intervals (CrIs) are shown by black vertical lines. The posterior means and CrIs are shown for the average disc width of female and male individuals caught within the corresponding location and season.

**Table S3. Stable isotope ratios of potential prey species of southern stingrays used in the Bayesian mixing models.** We used the values and the number digits after the decimal points as they were reported in the literature.

| Species | Group | $\delta^{13}C$ *[‰]* | $\delta^{15}N$ *[‰]* | Source |
|---|---|---|---|---|
| *Caridea* | Crustaceans | -13.76 | 6.17 | Tilley et al. 2013 |
| *Panulirus argus* | Crustaceans | -11.92 | 5.75 | Tilley et al. 2013 |
| Juvenile *Panulirus argus* | Crustaceans | -13.52 | 5.65 | Tilley et al. 2013 |
| *Bivalvia* | Molluscs | -12.03 | 4.38 | Tilley et al. 2013 |
| *Aliger gigas* | Molluscs | -12.38 | 3.37 | Tilley et al. 2013 |
| *Arenicola* | Annelids | -8.04 | 2.79 | Tilley et al. 2013 |
| *Brachyura spp.* | Crustaceans | -14.96 | 2.96 | Tilley et al. 2013 |
| *Aplysia dactylomela* | Molluscs | -18.56 | 4.0 | Tilley et al. 2013 |
| *Coryphopteris personatus* | Teleosts | -16.5 | 4.0 | Zhu et al. 2019 |
| *Coryphotpreus glaucofraenum* | Teleosts | -10.9 | 3.9 | Zhu et al. 2019 |
| *Elacantinus genie* | Teleosts | -12.6 | 8.9 | Zhu et al. 2019 |
| Unidentified polychaetes | Annelids | -23.7 | 6.7 | Hoopes et al. 2020 |
| Unidentified polychaetes | Annelids | -15.8 | 6.5 | Hoopes et al. 2020 |
| Unidentified *Bivalvia* | Molluscs | -9.7 | 3.5 | Hoopes et al. 2020 |
| *Pteria colymbus* | Molluscs | -13.6 | 5.0 | Hoopes et al. 2020 |
| *Aliger gigas* | Molluscs | -12.2 | 3.6 | Hoopes et al. 2020 |
| *Menippe mercenaria* | Crustaceans | -19.5 | 5.3 | Hoopes et al. 2020 |
| *Carcinus meanus* | Crustaceans | -12.7 | 8.5 | Hoopes et al. 2020 |
| *Callinectes sapidus* | Crustaceans | -15.7 | 9.0 | Hoopes et al. 2020 |
| *Panulirus argus* | Crustaceans | -12.0 | 5.7 | Hoopes et al. 2020 |
| *Litopenaeus setiferus* | Crustaceans | -20.6 | 10.1 | Hoopes et al. 2020 |
| *Alpheus heterochaelis* | Crustaceans | -22.7 | 6.6 | Hoopes et al. 2020 |
| *Lysmata wurdemanni* | Crustaceans | -20.6 | 6.9 | Hoopes et al. 2020 |
| *Penaeus spp.* | Crustaceans | -15.6 | 12.9 | Hoopes et al. 2020 |
| *Opsanus beta* | Teleosts | -20.8 | 7.7 | Hoopes et al. 2020 |
| *Bathygobius soporator* | Teleosts | -22.6 | 9.7 | Hoopes et al. 2020 |
| *Gobiosoma robustum* | Teleosts | -21.6 | 8.1 | Hoopes et al. 2020 |
| *Gobiosoma bosc* | Teleosts | -25.1 | 9.2 | Hoopes et al. 2020 |
| *Gobiosoma spp.* | Teleosts | -24.6 | 11.0 | Hoopes et al. 2020 |

| | | | | |
|---|---|---|---|---|
| *Lophogobius cyprinoides* | Teleosts | -24.4 | 9.5 | Hoopes et al. 2020 |
| *Lupinoblennius nicholsi* | Teleosts | -23.6 | 11.3 | Hoopes et al. 2020 |
| *Coryphopterus personatus* | Teleosts | -17.1 | 6.8 | Hoopes et al. 2020 |
| *Coryphopterus glaucofraenum* | Teleosts | -10.9 | 3.9 | Hoopes et al. 2020 |
| *Elacatinus genie* | Teleosts | -12.6 | 8.9 | Hoopes et al. 2020 |
| *Isopoda/amphipoda* | Crustaceans | -16.9 | 7.7 | Kieckbusch et al. 2004 |
| *Panopeus herbstii* | Crustaceans | -13.5 | 4.3 | Kieckbusch et al. 2004 |
| *Isognomon alatus* | Molluscs | -14.9 | 4.9 | Kieckbusch et al. 2004 |
| *Amphipoda* | Crustaceans | -17.5 | 5.9 | Kieckbusch et al. 2004 |
| *Microphrys bicornutus* | Crustaceans | -17.7 | 5.2 | Kieckbusch et al. 2004 |
| *Pachygrapsus transversus* | Crustaceans | -13.5 | 4.3 | Kieckbusch et al. 2004 |
| *Aplysia* | Molluscs | -13.1 | 4.5 | Kieckbusch et al. 2004 |
| *Callinectes similis* | Crustaceans | -16.8 | 8.0 | Kieckbusch et al. 2004 |
| *Pagurus* | Crustaceans | -18.1 | 3.1 | Kieckbusch et al. 2004 |
| *Atrina* | Molluscs | -14.9 | 3.3 | Kieckbusch et al. 2004 |

**Table S4. Metadata of all southern stingrays sampled for blood for the analysis of condition metrics in serum samples.** The PIT tag column contains the individual-specific number of the passive integrated transponder tag. “F” in the Sex column shows females individuals, and “M” male individuals. The DW columns contains the disc width measurement in cm. The units in which condition metrics are measured are shown in []. The last column contains the capture location of the individuals (see Fig. 1) with the abbreviations: “LAG” – lagoon, “BFH” – Bonefish Hole, and “SB” – South Bimini.

| PIT tag | Date | Sex | DW *[cm]* | Butyrate *[mmol/L]* | Glucose *[mg/dl]* | Lactate *[mmol/L]* | Osmolality *[mOsm]* | Location |
|---|---|---|---|---|---|---|---|---|
| 989001026313458 | 2018-08-05 | F | 93.3 | 0.5 | 33.0 | 1.2 | 1158.0 | SB |
| 989001026313498 | 2018-10-28 | F | 85.2 | 0.5 | 48.0 | 0.6 | 1145.0 | SB |
| 989001026313524 | 2018-11-03 | F | 93.1 | 0.5 | 43.0 | 2.3 | 1174.0 | SB |
| 989001004378368 | 2018-11-08 | F | 101.2 | 0.6 | 43.0 | 1.9 | 1163.0 | SB |
| 989001004605300 | 2018-11-08 | F | 94.4 | 0.6 | 25.0 | 2.4 | 1176.0 | SB |
| 989001006700493 | 2018-11-08 | F | 78.1 | 0.6 | 43.0 | 2.5 | 1168.0 | SB |
| 900236000107745 | 2018-11-15 | F | 98.8 | 0.7 | 38.0 | 1.9 | 1161.0 | SB |
| 989001026313302 | 2018-11-17 | F | 102.1 | 0.5 | 32.0 | 2.0 | 1136.0 | SB |
| 989001004605145 | 2018-11-17 | F | 90.1 | 0.5 | 45.0 | 1.3 | 1172.0 | SB |
| 989001006700520 | 2018-11-17 | F | 94.0 | 0.5 | 44.0 | 2.2 | 1171.0 | SB |
| 989001005924837 | 2018-11-17 | F | 104.8 | 0.5 | 36.0 | 2.0 | 1170.0 | SB |
| 989001026313143 | 2018-11-17 | F | 77.0 | 0.6 | 41.0 | 0.9 | 1158.0 | SB |
| 989001005924813 | 2018-11-17 | F | 65.2 | 0.4 | 42.0 | 1.1 | 1165.0 | SB |
| 989001026313413 | 2018-11-26 | F | 77.0 | 0.4 | 46.0 | 0.6 | 1135.0 | SB |
| 989001026313157 | 2018-11-26 | F | 84.0 | 0.4 | 71.0 | 1.6 | 1137.0 | SB |
| 989001026313537 | 2018-11-26 | F | 66.8 | 0.5 | 50.0 | 1.5 | 1152.0 | SB |
| 989001005924851 | 2018-11-26 | F | 79.4 | 0.5 | 46.0 | 0.5 | 1151.0 | SB |
| 900236000107745 | 2018-12-01 | F | 98.8 | 0.5 | 58.0 | 1.4 | 1112.0 | SB |
| 989001026313275 | 2018-12-01 | F | 91.1 | 0.7 | 47.0 | 0.3 | 1145.0 | SB |
| 989001026313219 | 2018-12-01 | F | 87.2 | 0.6 | 46.0 | 0.9 | 1087.0 | SB |
| 985121031809367 | 2018-12-01 | F | 96.0 | 0.6 | 50.0 | 1.2 | 1117.0 | SB |
| 989001026313529 | 2018-12-01 | F | 92.0 | 0.5 | 44.0 | 0.6 | 1138.0 | SB |
| 989001004605145 | 2018-12-10 | F | 93.0 | 0.6 | 58.0 | 2.4 | 1132.0 | SB |
| 989001006700810 | 2018-12-10 | F | 88.3 | 0.4 | 55.0 | 0.9 | 1134.0 | SB |
| 4C4A72220F | 2018-12-10 | F | 96.4 | 0.7 | 45.0 | 0.3 | 1131.0 | SB |

| | | | | | | | | |
|---|---|---|---|---|---|---|---|---|
| 989001026313492 | 2018-12-10 | F | 76.1 | 0.6 | 49.0 | 0.3 | 1108.0 | SB |
| 989001006700794 | 2019-01-25 | F | 95.0 | 0.5 | 52.0 | 1.8 | 1132.0 | SB |
| 985121031809367 | 2019-01-25 | F | 94.9 | 0.7 | 42.0 | 0.7 | 1136.0 | SB |
| 985121031809367 | 2019-01-25 | F | 94.9 | 0.7 | 42.0 | 0.7 | 1136.0 | SB |
| 989001026313278 | 2019-01-25 | M | 42.2 | 0.6 | 55.0 | 2.6 | 1166.0 | SB |
| 989001006700707 | 2019-01-25 | F | 67.1 | 0.5 | 49.0 | 0.5 | 1159.0 | SB |
| 989001006700847 | 2019-01-25 | F | 91.1 | 0.7 | 41.0 | 1.0 | 1125.0 | SB |
| 989001004378382 | 2019-02-04 | F | 82.4 | 0.6 | 37.0 | 0.6 | 1105.0 | SB |
| 989001004378382 | 2019-02-04 | F | 82.4 | 0.6 | 33.0 | 0.8 | 1144.0 | SB |
| 989001026313255 | 2019-02-06 | M | 50.0 | 0.4 | 49.0 | 1.3 | 1171.0 | SB |
| 989001006700847 | 2019-02-07 | F | 94.9 | 2.5 | 31.0 | 1.6 | 1152.0 | SB |
| 989001026313200 | 2019-02-07 | F | 91.8 | 0.5 | 45.0 | 0.6 | 1123.0 | SB |
| 989001004378515 | 2019-02-07 | F | 82.0 | 0.6 | 50.0 | 0.7 | 1137.0 | SB |
| 989001026313388 | 2019-03-16 | F | 85.8 | 0.5 | 52.0 | 2.5 | 1159.0 | BFH |
| 989001026313357 | 2019-03-16 | F | 87.4 | 0.6 | 47.0 | 0.7 | 1162.0 | BFH |
| 989001026313378 | 2019-03-16 | F | 82.0 | 0.5 | 47.0 | 0.5 | 1169.0 | BFH |
| 989001026313172 | 2019-03-16 | F | 95.6 | 0.5 | 39.0 | 2.4 | 1181.0 | BFH |
| 989001026313343 | 2019-03-17 | F | 74.0 | 0.5 | 50.0 | 0.3 | 1164.0 | BFH |
| 989001005924856 | 2019-03-17 | F | 74.6 | 0.4 | 30.0 | 0.4 | 1163.0 | BFH |
| 989001026313187 | 2019-03-25 | F | 108.4 | 0.4 | 40.0 | 0.6 | 1074.0 | LAG |
| 989001026313359 | 2019-03-25 | F | 92.0 | NA | 75.0 | 2.1 | 1133.0 | LAG |
| 989001026313370 | 2019-03-25 | M | 54.2 | 0.4 | 22.0 | 0.3 | 1100.0 | LAG |
| 989001026313402 | 2019-03-25 | F | 88.5 | 0.5 | 43.0 | 0.6 | 1106.0 | LAG |
| 989001026313371 | 2019-03-26 | F | 83.0 | 0.4 | 30.0 | 0.8 | 928.0 | SB |
| 989001005924975 | 2019-03-26 | F | 97.4 | 0.5 | 45.0 | 1.1 | 1100.0 | SB |
| 989001006700794 | 2019-03-26 | F | 97.0 | 0.5 | 47.0 | 1.2 | 1095.0 | SB |
| 985121031817057 | 2019-03-26 | F | 91.2 | 0.6 | 42.0 | 0.6 | 1092.0 | SB |
| NoTag | 2019-03-26 | F | 84.0 | 0.4 | 36.0 | 2.4 | 1117.0 | SB |
| 989001026313440 | 2019-03-26 | M | 48.8 | 0.6 | 49.0 | 1.8 | 1122.0 | SB |
| 989001026313197 | 2019-03-26 | F | 72.6 | 0.6 | 51.0 | 1.0 | 1029.0 | SB |
| 989001004378368 | 2019-04-20 | F | 94.0 | 0.6 | 54.0 | 0.9 | 1142.0 | SB |
| 989001026313194 | 2019-05-09 | F | 103.4 | 0.6 | 18.0 | 3.1 | 1167.0 | SB |
| 989001026313417 | 2019-05-09 | F | 93.9 | 0.5 | 45.0 | 2.3 | 1151.0 | BFH |
| 985121031803859 | 2019-05-09 | F | 88.3 | 0.5 | 42.0 | 1.1 | 1163.0 | BFH |

| | | | | | | | | |
|---|---|---|---|---|---|---|---|---|
| 989001026313525 | 2019-05-09 | F | 90.3 | 0.4 | 25.0 | 4.1 | 1130.0 | BFH |
| 989001026313152 | 2019-05-09 | F | 92.5 | 0.5 | 35.0 | 1.3 | 1163.0 | BFH |
| 989001026313486 | 2019-05-09 | F | 88.0 | 0.5 | 21.0 | 2.7 | 1170.0 | BFH |
| 900236000108035 | 2019-06-28 | F | 74.5 | 0.6 | 51.0 | 1.0 | 1116.0 | SB |
| 989001026313198 | 2019-07-01 | M | 45.0 | 0.6 | 56.0 | 1.1 | 1105.0 | SB |
| 989001026313493 | 2019-07-01 | F | 80.2 | 0.4 | 62.0 | 2.4 | 1089.0 | SB |
| 989001006700567 | 2019-07-01 | F | 86.5 | 0.5 | 59.0 | 2.5 | 1093.0 | SB |
| 989001026313243 | 2019-07-01 | F | 86.0 | 0.5 | 43.0 | 1.7 | 1110.0 | SB |
| 989001026313369 | 2019-07-09 | F | 92.0 | 0.6 | 22.0 | 3.1 | 1149.0 | BFH |
| 989001026313510 | 2019-07-09 | F | 89.0 | 0.5 | 21.0 | 3.6 | 1129.0 | BFH |
| 989001006700560 | 2019-07-09 | F | 92.6 | 0.5 | 58.5 | 2.5 | 1152.0 | BFH |
| 989001026313193 | 2019-07-09 | F | 72.5 | 0.6 | 24.0 | 3.2 | 1169.0 | BFH |
| 989001026313480 | 2019-07-20 | F | 69.5 | 0.4 | 29.0 | 3.8 | 1195.0 | LAG |
| 989001026313456 | 2019-07-20 | F | 98.0 | 0.4 | 35.0 | 2.3 | 1169.0 | LAG |
| 989001026313538 | 2019-07-20 | F | 82.0 | 0.4 | 16.0 | 4.4 | 1176.0 | SB |
| 989001026313468 | 2019-07-22 | F | 83.0 | 0.4 | 18.0 | 3.5 | 1231.5 | LAG |
| 989001026313512 | 2019-07-22 | F | 96.0 | 0.4 | 30.0 | 3.2 | 1203.0 | LAG |
| 989001026313449 | 2019-07-22 | F | 65.4 | 0.4 | 40.0 | 4.5 | 1254.0 | LAG |
| 989001026313464 | 2019-07-22 | F | 105.6 | 0.5 | 57.0 | 4.4 | 1178.0 | SB |
| 989001026313161 | 2019-08-13 | F | 56.4 | 0.5 | 61.0 | 1.6 | 1196.0 | SB |
| 989001026313442 | 2019-08-21 | M | 42.6 | 0.6 | 31.0 | 2.7 | 1154.0 | SB |
| 900236000107745 | 2019-08-21 | F | 93.5 | 0.6 | 23.0 | 3.9 | 1136.0 | SB |
| 989001004378459 | 2019-08-21 | F | 93.7 | 0.5 | 40.0 | 2.6 | 1247.0 | SB |
| 989001026313424 | 2019-09-15 | F | 80.2 | 0.5 | 39.0 | 4.0 | 1153.0 | LAG |
| 989001026313473 | 2019-09-15 | F | 79.5 | 0.5 | 25.0 | 3.3 | 1120.0 | LAG |
| 989001026313532 | 2019-09-15 | F | 88.0 | 0.4 | 67.0 | 5.4 | 1136.0 | LAG |
| 989001006700537 | 2019-09-15 | F | 60.0 | 0.4 | 51.0 | 2.8 | 1106.0 | LAG |
| 989001026313495 | 2019-09-15 | F | 76.5 | 0.5 | 20.0 | 3.5 | 1152.0 | LAG |
| 989001030278548 | 2019-09-19 | F | 78.0 | 0.4 | 40.0 | 1.3 | 1129.0 | BFH |
| 989001030278436 | 2019-09-19 | F | 70.8 | 0.4 | 53.0 | 0.6 | 1138.0 | BFH |
| 989001026313488 | 2019-09-24 | F | 54.2 | 0.5 | 51.0 | 0.6 | 1133.0 | SB |
| 989001005924875 | 2019-09-24 | F | 92.1 | 0.6 | 46.0 | 3.4 | 1133.0 | SB |
| 989001030278412 | 2019-10-04 | F | 90.4 | 0.5 | 39.0 | 2.7 | 1143.0 | BFH |
| 989001026313469 | 2019-10-04 | F | 101.0 | 0.4 | 27.0 | 4.4 | 1243.0 | BFH |

| | | | | | | | | |
|---|---|---|---|---|---|---|---|---|
| 989001030278440 | 2019-10-04 | F | 91.5 | 0.6 | 32.0 | 2.4 | 1156.0 | BFH |
| 989001030278261 | 2019-10-04 | F | 83.0 | 0.5 | 59.0 | 1.4 | 1183.0 | BFH |
| 989001030278485 | 2019-10-04 | F | 70.4 | 0.5 | 10.0 | 4.7 | 1171.0 | BFH |
| 989001030278490 | 2019-10-05 | F | 78.0 | 0.6 | 37.0 | 2.4 | 1283.0 | LAG |
| 989001030278456 | 2019-10-05 | F | 80.3 | 0.4 | 27.0 | 3.8 | 1101.0 | LAG |
| 989001030278533 | 2019-10-05 | F | 73.0 | 0.5 | 22.0 | 5.0 | 1219.0 | LAG |
| 989001030278420 | 2019-10-05 | F | 68.3 | 0.5 | 53.0 | 1.2 | 1160.0 | LAG |
| 989001030278298 | 2019-10-13 | M | 45.0 | 0.8 | 69.0 | 2.9 | 1197.0 | SB |
| 989001030278610 | 2019-10-21 | F | 64.0 | 0.5 | 60.0 | 1.6 | 1198.0 | SB |
| 989001026313461 | 2019-10-21 | M | 51.6 | 0.4 | 65.0 | 1.4 | 1131.0 | SB |
| 989001026313180 | 2019-11-01 | F | 77.0 | 0.4 | 36.0 | 1.4 | 1132.0 | LAG |
| 989001030278429 | 2019-11-01 | F | 90.5 | 0.4 | 10.0 | 4.8 | 1117.0 | LAG |
| 989001030278216 | 2019-11-01 | F | 85.7 | 0.5 | 25.0 | 4.5 | 1133.0 | LAG |
| 989001030278291 | 2019-11-01 | F | 88.5 | 0.4 | 36.0 | 2.5 | 1244.0 | LAG |
| 989001030278233 | 2019-11-01 | F | 91.7 | 0.4 | 14.0 | 4.7 | 1130.0 | LAG |
| 989001030278549 | 2019-11-01 | F | 73.2 | 0.4 | 59.0 | 2.3 | 1150.0 | LAG |
| 989001030278271 | 2019-11-01 | F | 78.4 | 0.5 | 49.0 | 1.0 | 1135.0 | LAG |
| 989001030278434 | 2019-11-01 | M | 51.9 | 0.4 | 37.0 | 1.5 | 1152.0 | LAG |
| 989001030278304 | 2019-11-01 | F | 72.5 | 0.4 | 22.0 | 3.5 | 1132.0 | LAG |
| 989001030278237 | 2019-11-01 | F | 78.5 | 0.7 | 56.0 | 0.5 | 1121.0 | LAG |
| 989001030278310 | 2019-11-12 | F | 52.0 | 2.5 | 63.0 | 2.1 | 1163.0 | SB |
| 989001030278567 | 2019-11-12 | F | 71.0 | 0.6 | 50.0 | 1.5 | 1151.0 | SB |
| 989001030278219 | 2019-11-12 | F | 63.5 | 0.7 | 47.0 | 1.1 | 1134.0 | SB |
| 989001030278446 | 2019-11-12 | M | 53.0 | 0.8 | 51.0 | 1.6 | 1163.0 | SB |
| 989001006700707 | 2019-11-12 | F | 71.2 | 2.5 | 40.0 | 1.2 | 1151.0 | SB |
| 989001030278551 | 2019-11-25 | F | 86.1 | 0.4 | 10.0 | 4.0 | 1138.0 | BFH |
| 989001030278599 | 2019-11-25 | F | 92.7 | 0.6 | 34.0 | 2.5 | 1147.0 | BFH |
| 989001030278545 | 2019-11-25 | F | 99.5 | 0.8 | 22.0 | 2.9 | 1141.0 | BFH |
| 989001030278265 | 2019-11-27 | F | 84.8 | 0.5 | 40.0 | 0.6 | 1130.0 | BFH |
| 989001030278562 | 2019-11-27 | F | 97.9 | 0.5 | 51.0 | 2.4 | 1144.0 | BFH |
| 989001030278435 | 2019-11-27 | F | 89.0 | 0.4 | 30.0 | 1.2 | 1146.0 | BFH |

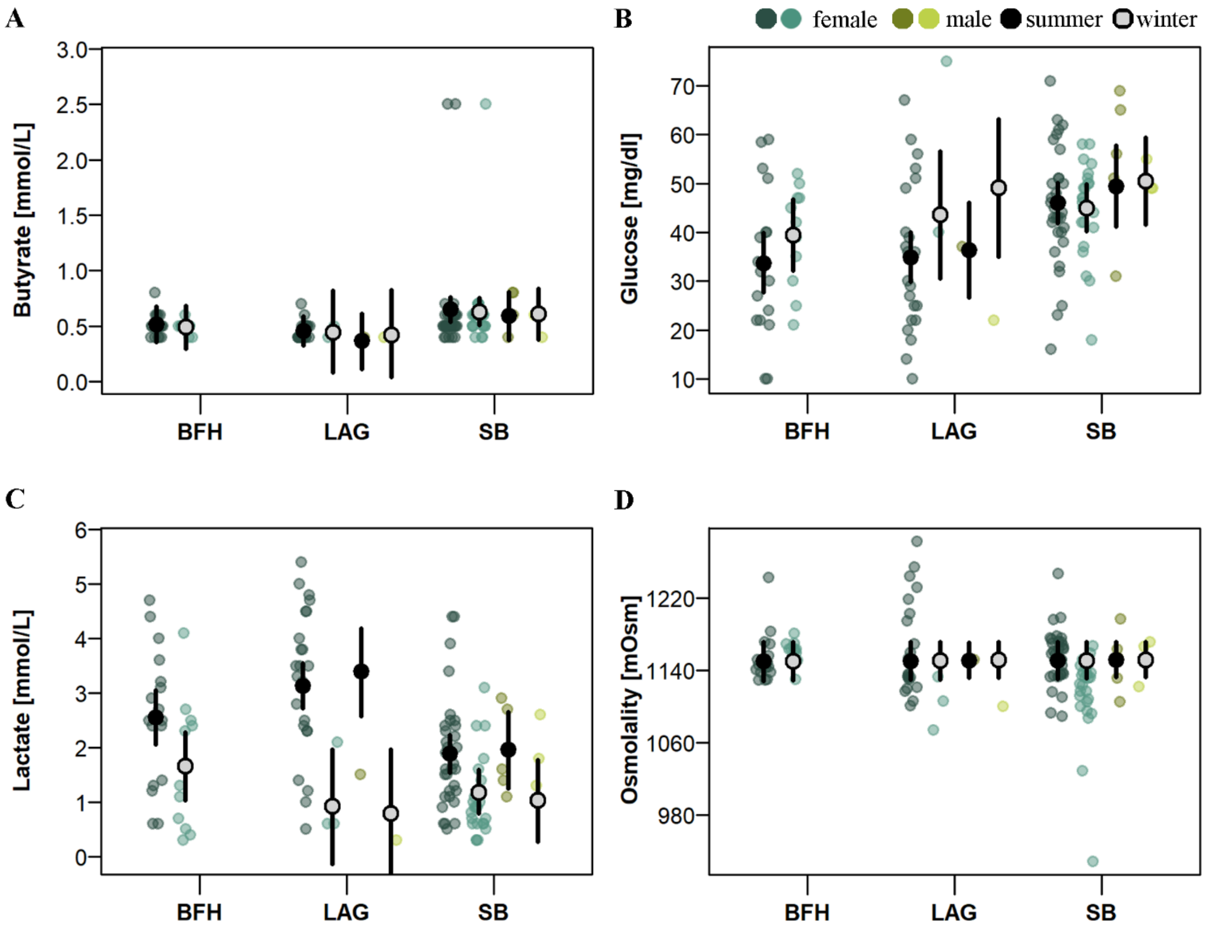


**Fig. S4. Effects of sex, location, and season on concentrations of (A) butyrates, (B) glucose, (C) lactate, and (D) osmolality in serum samples.** Coloured points show the raw data points by sex and season. Raw data points for female and male individuals are shown for the two seasons summer (darker colours) and winter (lighter colours). Capture locations are abbreviated as "LAG" – lagoon, "BFH" – Bonefish Hole, and "SB" – South Bimini. Posterior means simulated from the statistical models are shown by black (summer) and grey points (winter) and 95 % credible intervals (CrIs) are shown by black vertical lines. The posterior means and CrIs are shown for the average disc width of female and male individuals caught within the corresponding location and season.

*Supplementary references*


Hoopes, L. A., Clauss, T. M., Browning, N. E., Delaune, A. J., Wetherbee, B. M., Shivji, M., Harvey, J. C., & Harvey, G. C. M. (2020). Seasonal patterns in stable isotope and fatty acid profiles of southern stingrays (*Hypanus americana*) at Stingray City Sandbar, Grand Cayman. *Scientific Reports*, *10*, 19753. https://doi.org/10.1038/s41598-020-76858-w

Kieckbusch, D. K., Koch, M. S., Serafy, J. E., & Anderson, W. T. (2004). Trophic linkages among primary producers and consumers in fringing mangroves of subtropical lagoons. *Bulletin of Marine Science*, *74*(2), 271–285.

Tilley, A., López-Angarita, J., & Turner, J. R. (2013). Diet Reconstruction and Resource Partitioning of a Caribbean Marine Mesopredator Using Stable Isotope Bayesian Modelling. *PLoS ONE*, *8*(11), e79560. https://doi.org/10.1371/journal.pone.0079560

Zhu, Y., Newman, S. P., Reid, W. D. K., & Polunin, N. V. C. (2019). Fish stable isotope community structure of a Bahamian coral reef. *Marine Biology*, *166*(12), 160. https://doi.org/10.1007/s00227-019-3599-9
